# Critical Role for the Unique N-Terminus of Chlamydial MreB in Directing Its Membrane Association and Interaction with Elements of the Divisome

**DOI:** 10.1101/697953

**Authors:** Junghoon Lee, John V. Cox, Scot P. Ouellette

## Abstract

*Chlamydiae* lack the conserved central coordinator protein of cell division FtsZ, a tubulin-like homolog. Current evidence indicates *Chlamydia* uses the actin-like homolog, MreB, to substitute for the role of FtsZ. Interestingly, we observed MreB as a ring at the septum in dividing cells of *Chlamydia*. We hypothesize that MreB, to substitute for FtsZ in *Chlamydia*, must possess unique properties compared to canonical MreB orthologs. Sequence differences between chlamydial MreB and orthologs in other bacteria revealed that chlamydial MreB possesses an extended N-terminal region and the conserved amphipathic helix found in other bacterial MreBs. The extended N-terminal region was sufficient to restore the localization of a truncated *E. coli* MreB mutant lacking its amphipathic helix to the membrane and was crucial for interactions with cell division components RodZ and FtsK, though the region was not required for homotypic interactions. Importantly, the N-terminal region was sufficient to direct GFP to the membrane when expressed in *Chlamydia*. A mutant N-terminal region with reduced amphipathicity was unable to perform these functions. From these data, the extended N-terminal region of chlamydial MreB is critical for localization and interactions of this protein. Our data provide mechanistic support for chlamydial MreB to serve as a substitute for FtsZ.

**Importance:** *Chlamydia trachomatis* is an obligate intracellular pathogen, causing sexual transmitted diseases and trachoma. Studying chlamydial physiology, especially its cell division mechanism, is important for developing novel therapeutic strategies for the treatment of these diseases. Since chlamydial cell division has unique features, including a polarized cell division process independent of FtsZ, a canonical cell division coordinator, studying the subject is helpful for understanding undefined aspects of chlamydial growth. In this study, we characterized MreB, a substitute for FtsZ, as a cell division coordinator. It forms a filamentous ring at the septum, like FtsZ in *E. coli*. We show that the localization of MreB is dependent upon the amphipathic nature of its extended N-terminus. Furthermore, this region is crucial for its interaction with other proteins involved in cell division. Given these results, chlamydial MreB may function as a scaffold for cell divisome proteins at the septum and regulate cell division in this organism.

## Introduction

Bacteria within the genus *Chlamydia* are obligate intracellular pathogens that cause diverse diseases in humans and animals. These Gram-negative cocci differentiate between two morphologically and functionally distinct cellular forms during their developmental cycle: the elementary body (EB) and the reticulate body (RB)(1). The EB is the infectious but non-dividing form whereas the RB is the dividing but non-infectious form. After entering the host cell, *Chlamydia* remains within a membrane-bound parasitic organelle, termed an inclusion(2). RBs undergo multiple rounds of cell division until they engage a secondary differentiation program and convert to EBs, which are subsequently released from the host cell to propagate the infection.

In evolving to obligate intracellular dependence, *Chlamydia* has significantly reduced its genome, eliminating *ftsZ*(3), the conserved tubulin-like cell division coordinator of binary fission(4). However, *Chlamydia* encodes rod-shape determining proteins associated with peptidoglycan synthesis in the lateral cell wall of bacilli. That *Chlamydia* has retained these genes is unusual since these are coccoid bacteria and only synthesize peptidoglycan during division(5). The absence of FtsZ in chlamydiae suggests they may not divide by binary fission. Recent work from our labs supports this. We employed both live-and fixed-cell imaging to localize various markers, including a division protein (FtsQ(6)), during EB-to-RB differentiation and the first division of the RB. Interestingly, we observed that chlamydiae are highly polarized throughout this time and that division itself occurs via a polarized process(7). This polarized budding process remains the primary mode of division even at later times during the developmental cycle (Cox et al., *in prep*).

In 2012, we hypothesized and presented evidence that *Chlamydia* co-opted the rod-shape determining protein MreB to function as the central coordinator of division(8). To perform this function, we hypothesize that chlamydial MreB possesses unique properties. Many MreB homologs encode an N-terminal amphipathic helix that facilitates MreB association with the inner membrane(9). At the membrane it engages peptidoglycan machinery while forming short moving filaments(10–12). Here, we investigated the unique structural features of chlamydial MreB that allow it to substitute for FtsZ. Chlamydial MreB forms rings at the division site, and bioinformatics analysis indicates that chlamydial MreB encodes an extended N-terminal region (amino acids 1-23: aa1-23) in addition to the conserved amphipathic helix. The aa1-23 residues also encode predicted amphipathicity. We confirmed that the conserved amphipathic helix, but not aa1-23, is important for directing membrane-localization of GFP in *E. coli*. Nevertheless, aa1-23 were sufficient to direct GFP to the membrane in *Chlamydia*, and this localization was abolished when mutations were introduced in aa1-23 to reduce its amphipathicity. We determined the N-terminal region of chlamydial MreB is crucial for its interactions with cell division proteins but not its ability to form homo-oligomers. Our data indicate the N-terminal region of chlamydial MreB is important for membrane localization and interaction with the divisome. These properties allow it to function in a manner distinct from other MreB orthologs and regulate the cell division process in *Chlamydia.*

## Results

### Chlamydial MreB_6xH localizes as a ring at the division site

Given our hypothesis for the role of chlamydial MreB in directing polarized division, we examined its localization during the first division of an RB. We were unable to detect endogenous MreB with the antibody previously used to image the distribution of MreB in *Chlamydia*(13). We then attempted to image wild-type and various truncation mutants of MreB using a GFP sandwich (SW) fusion strategy that, in *E. coli*, was reported to be functional(14) (N-or C-terminal fluorescent protein fusions with MreB display an artifactual localization(15)). However, when we inducibly expressed MreB_GFP_SW_ in *C. trachomatis* L2, bacterial growth was significantly inhibited even with low levels of expression, as observed by the presence of a small number of enlarged bacteria (Suppl. Fig. 1). More importantly, the fusion protein did not localize to the membrane as reported (Suppl. Fig. 1; (13)). The reasons for this are not clear. We, therefore, opted to construct an MreB fusion with a small C-terminal hexahistidine tag (MreB_6xH). The human epithelial cell line, HeLa, was infected with a transformant of *C. trachomatis* L2 carrying a plasmid to inducibly express MreB_6xH. At 6 hours post-infection (hpi), expression of MreB_6xH was induced with anhydrotetracycline (aTc), and cells were fixed at 10.5 hpi, a time when the nascent RB has begun its first division(7). As seen in Figure 1, we observed using super-resolution structured illumination microscopy (SIM) the polar localization of the major outer membrane protein (MOMP) in the budding daughter cell as previously reported(7). Consistent with its proposed role in division(8), MreB_6xH was localized in a band-like structure across the septum and not in puncta as previously reported(13, 16). Interestingly, when 3D reconstructions were assembled from the SIM images, we observed that MreB_6xH formed ring-like structures at the septum with areas of more intense signal (Fig. 1 arrowheads). These areas of more intense signal may correspond to the puncta previously observed (13, 16).

**Figure 1.**
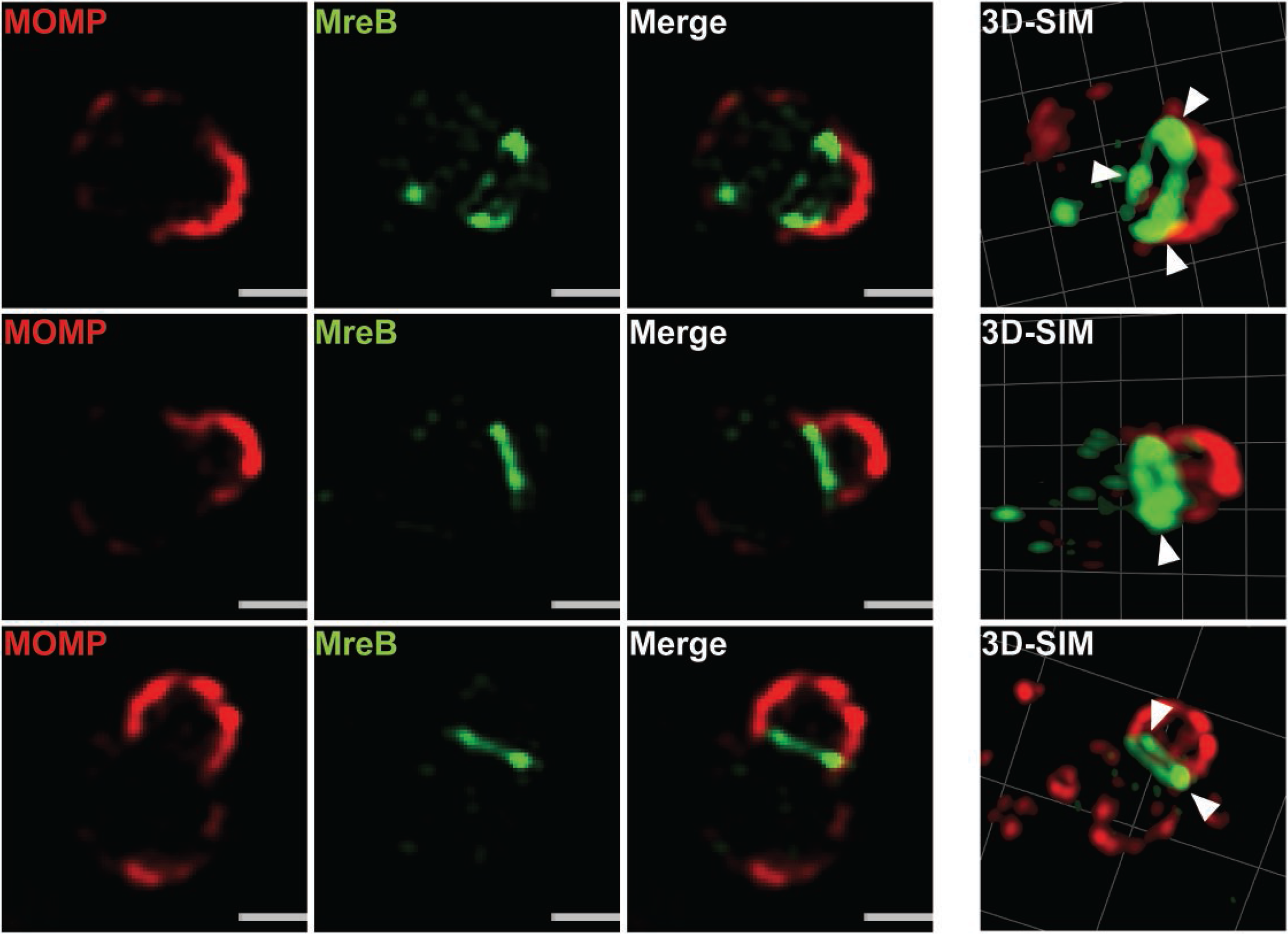
Localization of chlamydial MreB_6xH in *C. trachomatis* using structured illumination microscopy (SIM). *C. trachomatis* without plasmid (-pL2) was transformed with an anhydrotetracycline (aTc)-inducible vector encoding chlamydial MreB with a six-histidine (6xH) tag at the C-terminus. HeLa cells were infected with this strain and chlamydial MreB_6xH expression was induced with 10 nM aTc at 6 hpi. At 10.5 hpi, the infected cells were fixed (3.2% Formaldehyde, 0.022% Glutaraldehyde in PBS) for 2 min and permeabilized with 90% methanol (MeOH) for 1 min. The sample was stained for major outer membrane protein (MOMP; red) and chlamydial MreB (green). Three representative images are displayed. The arrowheads indicate regions of more intense fluorescence. Structured illumination microscope (SIM) images were acquired on a Zeiss ELYRA PS.1 super-resolution microscope. The scale bar = 0.5 μm.

### Chlamydial MreB encodes an amphipathic helix and an extended N-terminal region conserved in *Chlamydia*

In many Gram-negative bacteria, there is an amphipathic helix at the N-terminus of MreB, and the helix is important for the membrane localization of MreB(9). This localization is crucial for the role of MreB in directing PG synthesis during lateral cell wall growth(9, 17). To confirm whether chlamydial MreB also has these features, we predicted its secondary structure and amphipathic regions using bioinformatics tools (Fig. 2). The alignment data show that chlamydial MreB encodes an extended N-terminal region (amino acids 1-23: aa1-23), which other Gram-negative bacteria lack (Fig. 2A). Importantly, this extended N-terminal region is conserved amongst *Chlamydia* (Suppl. Fig. 2). Following this region, there is a predicted amphipathic helix, from R24 to F29 (Fig. 2B), that aligns with the amphipathic helices of the other bacteria (Fig. 2A). A high amphipathic score (A-score) extends to residue F32 in chlamydial MreB. Interestingly, aa1-23 also has a high A-score though the region was not predicted to form an amphipathic helix (Fig. 2B). Similar results were seen with the more divergent *Waddlia* MreB homolog (Suppl. Fig. 2B). Nevertheless, performing a helical wheel analysis with 8 amino acid windows revealed that each window shows amphipathicity (Fig. 2C). Based on these observations, we conclude that the extended N-terminal region of chlamydial MreB may possess membrane-associating properties.

**Figure 2.**
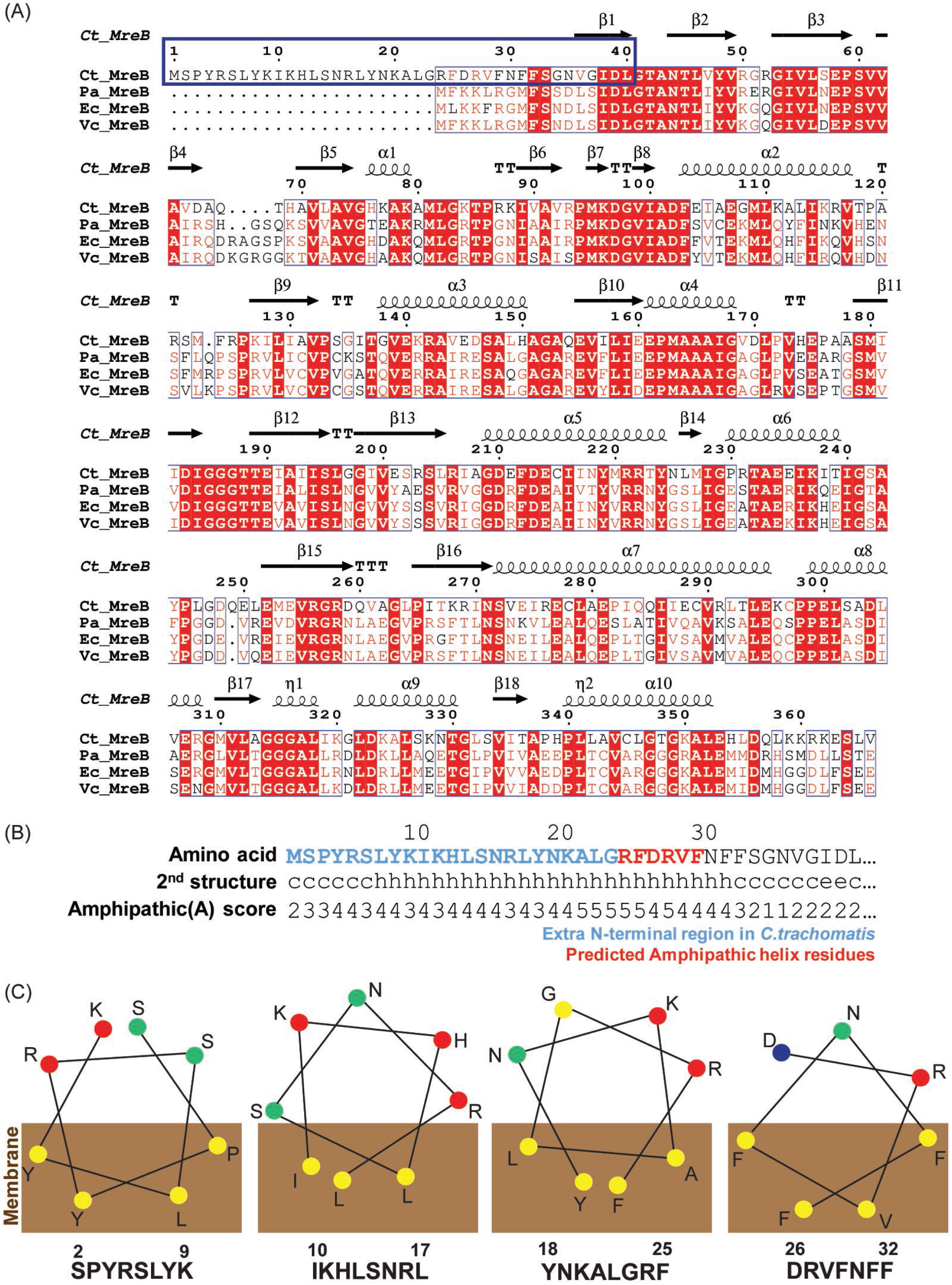
Bioinformatics analysis of chlamydial MreB. Chlamydial MreB has unique structural features based on bioinformatics analyses. (A) Protein sequence alignment of chlamydial and other bacterial MreBs. The blue box represents the extended N-terminus and predicted amphipathic helix of chlamydial MreB. The alignment was performed with Clustal Omega (https://www.ebi.ac.uk/Tools/msa/clustalo/) and represented with ESPript 3.0 (http://espript.ibcp.fr). (B) The amphipathic score and the predicted secondary structure of the N-terminus of chlamydial MreB. The prediction of amphipathicity is performed by using AMPHIPASEEK. The blue and red residues represent the extended N-terminus and predicted amphipathic helix, respectively. (C) The amphipathicity of the fragments of the N-terminal region of chlamydial MreB. These amphipathic structures were predicted by an online helical wheel program, named Helixator (http://www.tcdb.org/progs/helical_wheel.php). The circles represent the indicated amino acid residues. (yellow circle: non-polar residue; green circle: uncharged polar residue; red circle: basic polar residue; blue circle: acidic polar residue).

### Chlamydial MreB is unable to complement an *mreB*-deficient mutant of *E. coli*

The bioinformatics data suggested that chlamydial MreB encodes unique structural features absent in other bacterial MreBs (Fig. 2A). To assess the activities associated with these unique structural elements, we introduced chlamydial MreB into an *mreB*-deficient mutant of *E. coli.* As *mreB* is an essential gene in *E. coli*, Bendezu et al. made the strain P2733, which has a deletion in the *mreB* gene and is conditionally viable by the overexpression of the *ftsQAZ* operon(18). This strain has been used for complementation assays and grows as coccoid bacteria, since MreB is necessary for the maintenance of rod-shape(18). To test whether chlamydial MreB (CtrMreB) is capable of complementing the *mreB* deficient mutant, P2733 was transformed with an arabinose-inducible vector encoding chlamydial *mreB*. Cell shape was compared after inducing MreB expression in comparison to an empty vector negative control and an *E. coli* MreB (EcMreB) positive control (Suppl. Fig. 3A&B). When EcMreB is expressed from an arabinose-inducible promoter in the P2733 strain, the *E. coli* cells begin to adopt a rod shape, consistent with other reports(14, 18, 19). When chlamydial MreB was induced, the morphology of P2733 was unchanged compared to uninduced or empty vector controls, indicating that chlamydial MreB does not complement the *mreB*-deficient mutant of *E. coli* (Suppl. Fig. 3C). To determine whether the inability of chlamydial MreB to complement was due to its extended N-terminal region, we performed the same experiment with ΔN22 chlamydial MreB, which more closely resembles *E. coli* MreB in size and characteristics at the N-terminus. We observed that ΔN22 chlamydial MreB also does not complement the *mreB* deficient *E. coli*, suggesting that the extended N-terminal region is not the only reason for the failure to complement (Suppl. Fig. 3D – see also Discussion). Chlamydial MreB expression was confirmed in these experiments by western blotting (Suppl. Fig. 3E).

### The amphipathic helix, but not the extended N-terminal region, of chlamydial MreB is sufficient to direct GFP to the membrane in *E. coli*

We hypothesized that, because of their amphipathicity, the unique sequence elements at the N-terminus of chlamydial MreB could direct the membrane localization of MreB (Fig. 1). To test this, we performed a series of localization studies in *E. coli* using the N-terminal regions of chlamydial MreB (CtrMreB) fused to GFP. A similar strategy was used to demonstrate that two copies of the amphipathic helix of *E. coli* MreB (EcMreB) were sufficient to direct GFP to the membrane(9). We made various fusions in which either one or two copies of portions of the N-terminal region of chlamydial MreB were fused to GFP (illustrated in Fig. 3A). We then observed the localization of these proteins in both wild-type (MG1655) and MreB-deficient (P2733) *E. coli* (Fig. 3B and Suppl. Fig. 4). We observed that a single copy of aa1-32 (MreB_1-32_), encoding all high A-score residues, was sufficient to direct GFP to the membrane. Interestingly, aa1-28 (MreB_1-28_), encoding both the predicted amphipathic helix and the extended N-terminus, was also sufficient to direct GFP to the membrane, but, in wild-type cells, its localization was primarily restricted to the poles (Fig. 3C). Consistent with what has been observed in *E. coli*, two copies of aa23-32 (MreB_23-32_), encoding the predicted amphipathic helix, directed GFP to the membrane. Contrary to our prediction, one or two copies of aa1-23 (MreB_1-23_) was not able to direct GFP to the membrane even though the region has a high A-score (Fig. 2B).

**Figure 3.**
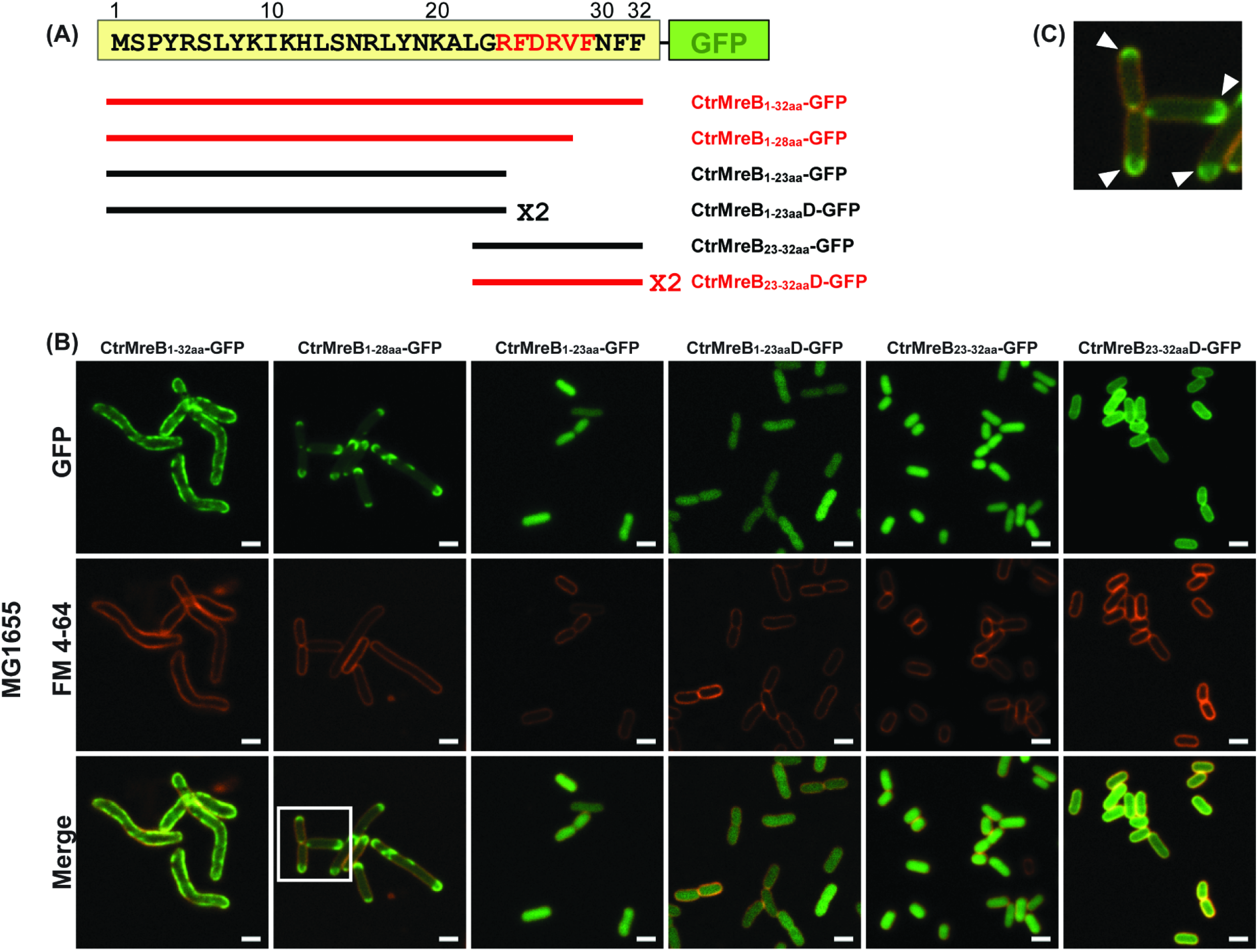
Localization in *E. coli* of various N-terminal regions of chlamydial MreB fused to GFP. Wild-type MG1655 *E. coli* was transformed with the arabinose-inducible vectors encoding GFP fusions with various N-terminal regions of chlamydial MreB (A-C). Stationary phase cultures strains were diluted 1:100 in LB media containing 34 μg/ml chloramphenicol and cultured for 2 h. The cells were then induced or not with 0.05% (w/v) arabinose and cultured for 2 h more. To stain the membrane, SynaptoRed-C2 (FM4-64) was added to the samples at a final concentration of 1.5 μM. After 15 min, 4 μL of each culture was spotted under a 1% LB agar pad and covered with a coverslip. The images were acquired on a Zeiss Imager.Z2 equipped with an Apotome2 using a 100X objective. (A) The structure of the various N-terminal MreB-GFP fusion peptides. In the boxed region containing the N-terminus of chlamydial MreB; red residues represent the predicted amphipathic helix residues. Red constructs show the membrane localization and black constructs show cytosolic localization. (B) The localization of various N-terminal regions of chlamydial MreB fused to GFP in MG1655. Note the polar membrane localization of the MreB_1-28aa_-GFP. Scale bar = 2 μm. (C) A zoomed image of MreB_1-28aa_-GFP localization represented by the box in (B). The arrowheads represent the polar membrane localization of the MreB_1-28aa_-GFP. Scale bar = 0.5 μm.

### The extended N-terminal region of chlamydial MreB is sufficient to direct GFP to the membrane in *C. trachomatis* L2

We next asked the question whether aa1-23, alone or in combination with the predicted amphipathic region (aa24-28), could direct GFP to the membrane in *Chlamydia*. The MreB_1-23aa__GFP or MreB_1-28aa__GFP fusions were moved to an inducible chlamydial expression plasmid and transformed into *C. trachomatis* L2. These transformants were then used to infect HeLa cells, and construct expression was induced at either 6 or 16 hpi. Infected cells were fixed and imaged at 10.5 or 20 hpi, respectively (Fig. 4). GFP alone is a cytosolic protein when expressed in chlamydiae (Fig. 4A). For both fusion proteins, GFP fluorescence was observed at membrane sites (Fig. 4B&C with arrowheads in 4B indicating individual bacteria with membrane localization of GFP) at both time points examined.

**Figure 4.**
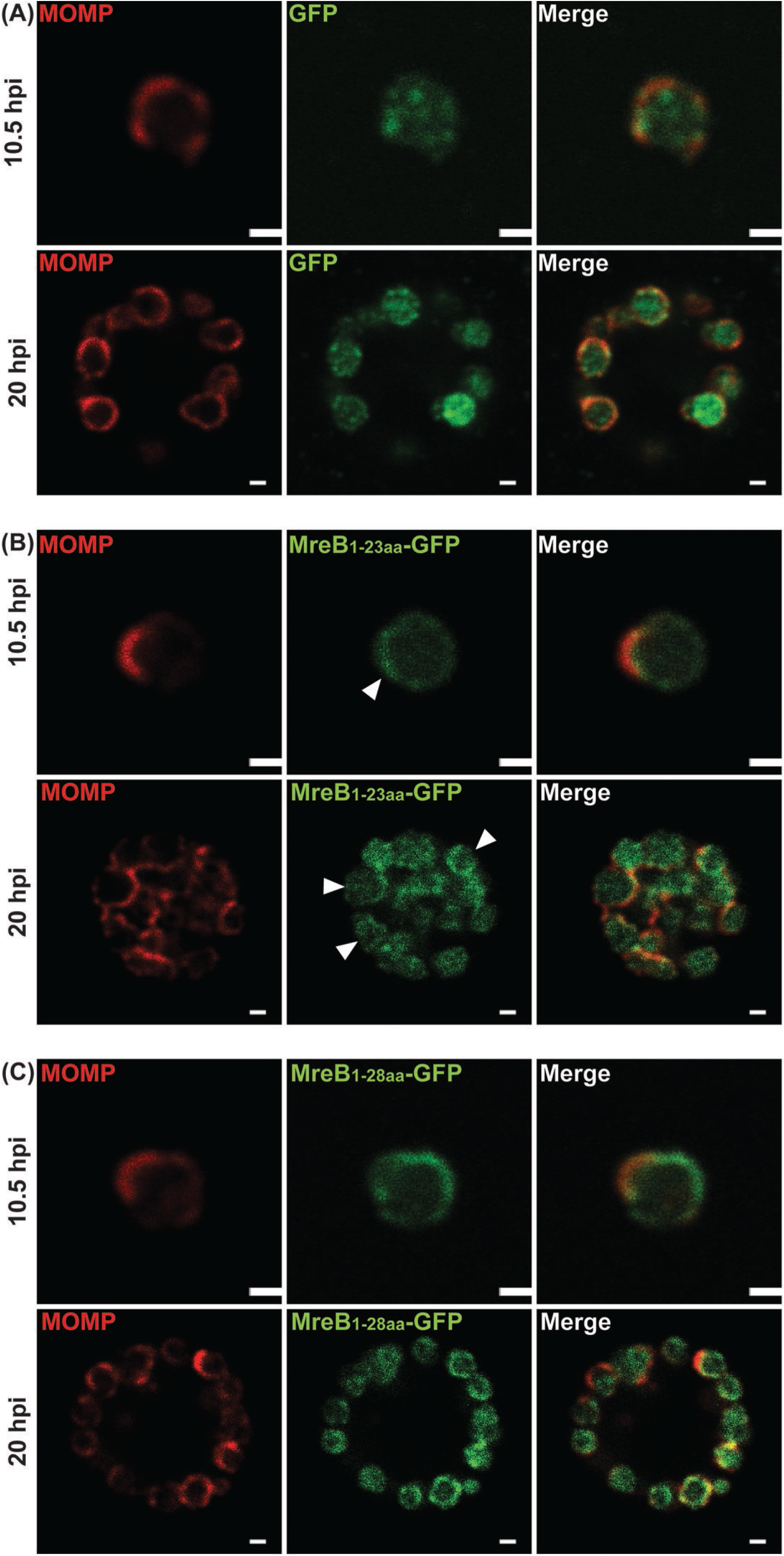
Localization in *C. trachomatis* of various N-terminal regions of chlamydial MreB fused to GFP. *C. trachomatis* serovar L2 without plasmid (-pL2) was transformed with anhydrotetracycline (aTc)-inducible vectors encoding (A) GFP, (B) chlamydial MreB1-23aa-GFP fusion peptide, or (C) chlamydial MreB_1-28aa_-GFP fusion peptide. HeLa cells were infected with the indicated strains, and expression of the GFP fusions was induced at 6 hpi or 16 hpi with 10 nM aTc. At 10.5 hpi or 20 hpi, the samples were fixed (3.2% Formaldehyde, 0.022% Glutaraldehyde in 1X PBS) for 2 min and permeabilized with 90% methanol (MeOH) for 1 min. These samples were stained for major outer membrane protein (MOMP; red) with GFP imaged in green. The arrowheads represent the MreB_1-23aa_-GFP localized at the membrane. Images were acquired on a Zeiss LSM 800 confocal microscope with 63X objective. Scale bar = 0.5 μm (10.5 hpi) or 1 μm (20 hpi).

Importantly, the aa1-23_GFP and aa1-28_GFP localization profiles were distinct from full-length MreB (Fig. 1, Suppl. Fig. 5), suggesting their localization profiles are not dependent on interactions with endogenous (i.e. chromosomally-encoded) MreB. To test this, we used the bacterial adenylate cyclase-based two hybrid (BACTH) assay, which is based on the reconstitution of enzyme activity by two interacting proteins that bring catalytic adenylate cyclase fragments (T25 and T18) into close proximity. We performed BACTH assays with full-length MreB and aa1-32 of MreB, and observed no interaction (data not shown). To further test a role for interactions with endogenous MreB, we expressed the GFP fusion proteins or MreB_6xH and then treated the cultures with A22, an MreB-specific antibiotic, to depolymerize MreB. Under these conditions, we observed the localization of the GFP fusion proteins at the membrane whereas the membrane-associated MreB_6xH was significantly reduced (Suppl. Fig. 5). Based on these data, we conclude that the membrane localization of the GFP fusion proteins is caused by the amphipathic nature of the N-terminus and not its interaction with endogenous MreB.

The N-terminal region of MreB from *C. trachomatis* has two leucine residues (L13 and L22) encoded by TTG. As bacteria can use UUG and GUG as alternative start codons (20), we performed a similar series of experiments using the N-terminal region of MreB from *C. suis*, which does not use UUG codons for its homologous leucine residues. In all cases tested, the N-terminal residues of *C. suis* MreB (fused to GFP) exhibited the same localization patterns as those observed for the *C. trachomatis* N-terminal MreB_GFP fusions (Suppl. Fig. 6). Based on these data, we conclude that it is unlikely the MreB of *C. trachomatis* bypasses the predicted AUG start codon in favor of downstream alternative start codons.

To further examine whether the membrane localization of aa1-23_GFP and aa1-28_GFP were dependent on their amphipathicity, we created the mutations L7K, L22R, and F25K. These mutations were predicted to diminish the amphipathicity of this region (Fig. 5A). The mutations within the N-terminal region were incorporated into the aa1-23_GFP and aa1-28_GFP fusion constructs, transformed into *C. trachomatis* L2, and inducibly expressed. As demonstrated in Figure 5 (B&C), we observed the cytosolic localization of the fusion peptides. Therefore, from the combination of these data, we conclude that the amphipathic properties of the extended N-terminal region of chlamydial MreB are sufficient to direct GFP to the membrane in *Chlamydia*.

**Figure 5.**
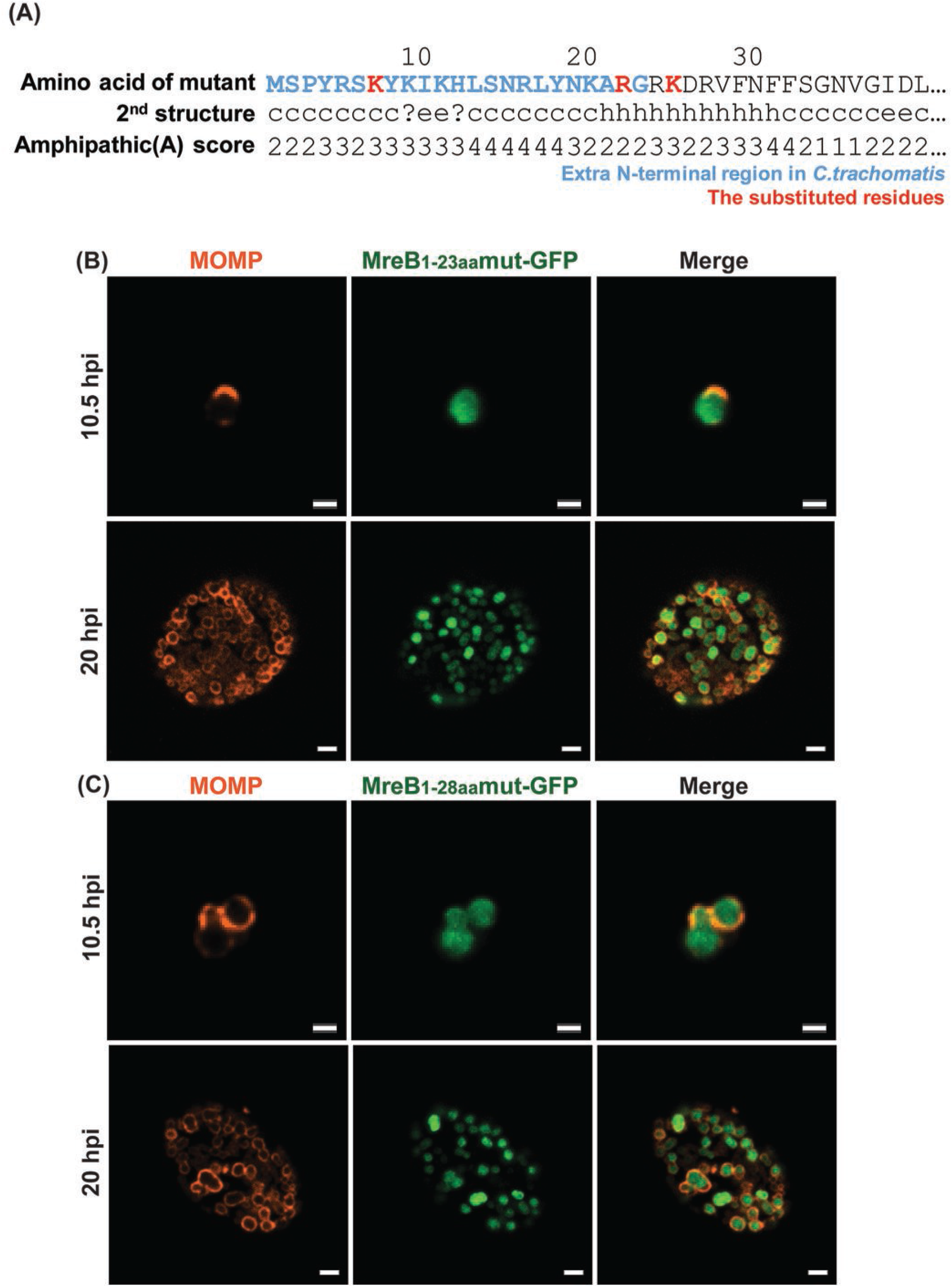
Localization in *C. trachomatis* of various mutated N-terminal regions of chlamydial MreB fused to GFP. (A) Mutations were introduced into the N-terminal region of chlamydial MreB and modeled for their effect on amphipathicity. L7K, L22R, and F25K (red residues) mutations disrupted the predicted amphipathicity (see Figure 2 for comparison). (B) *C. trachomatis* serovar L2 without plasmid (-pL2) was transformed with anhydrotetracycline (aTc)-inducible vectors encoding the mutated chlamydial MreB_1-23aa_- GFP or MreB_1-28aa_-GFP fusion peptide. HeLa cells were infected with the indicated strains, and expression of the GFP fusions was induced at 6 hpi or 16 hpi with 10 nM aTc. At 10.5 hpi or 20 hpi, the samples were fixed (3.2% Formaldehyde, 0.022% Glutaraldehyde in 1X PBS) for 2 min and permeabilized with 90% methanol (MeOH) for 1 min. These samples were stained for major outer membrane protein (MOMP; red) with GFP imaged in green. Images were acquired on a Zeiss Imager.Z2 equipped with an Apotome2 using a 100X objective. Scale bar = 1 μm (10.5 hpi) or 2 μm (20 hpi).

### The extended N-terminal region is dispensable for MreB homo-oligomerization but essential for interactions with other division proteins

We next asked whether the N-terminal region is necessary for MreB homo-oligomerization. In *E. coli*, MreB interacts with itself to form short filaments(10). In addition, since MreB participates in both cell shape determination and cell division, it interacts with diverse proteins related to cell division and peptidoglycan synthesis(14, 19, 21). These features are shared with chlamydial MreB, which interacts with itself and with several proteins such as the cytoskeletal protein RodZ and the cell division components FtsK and FtsQ(6, 8, 22). Based on these previous reports, we hypothesized that the extended N-terminal region is important for these interactions. To test this, we used the bacterial adenylate cyclase-based two hybrid (BACTH) assay. We designed BACTH constructs encoding truncated MreB mutants and then performed the interaction assays (Fig. 6). The N-terminal truncations of MreB interacted with full-length MreB (Fig. 6A and summarized in B), providing a potential explanation for the localization data of MreB_6xH truncations in *Chlamydia* (Suppl. Fig. 7). However, these truncated MreBs did not interact with RodZ, full-length FtsK, or the N-terminal region of FtsK lacking its ATPase domain whereas full-length MreB did interact with these constructs (Fig. 6C and summarized in D)(8, 22). From these data, we conclude that the extended N-terminal region is dispensable for MreB homo-oligomerization but is critical for interacting with other proteins involved in the cell division process.

**Figure 6.**
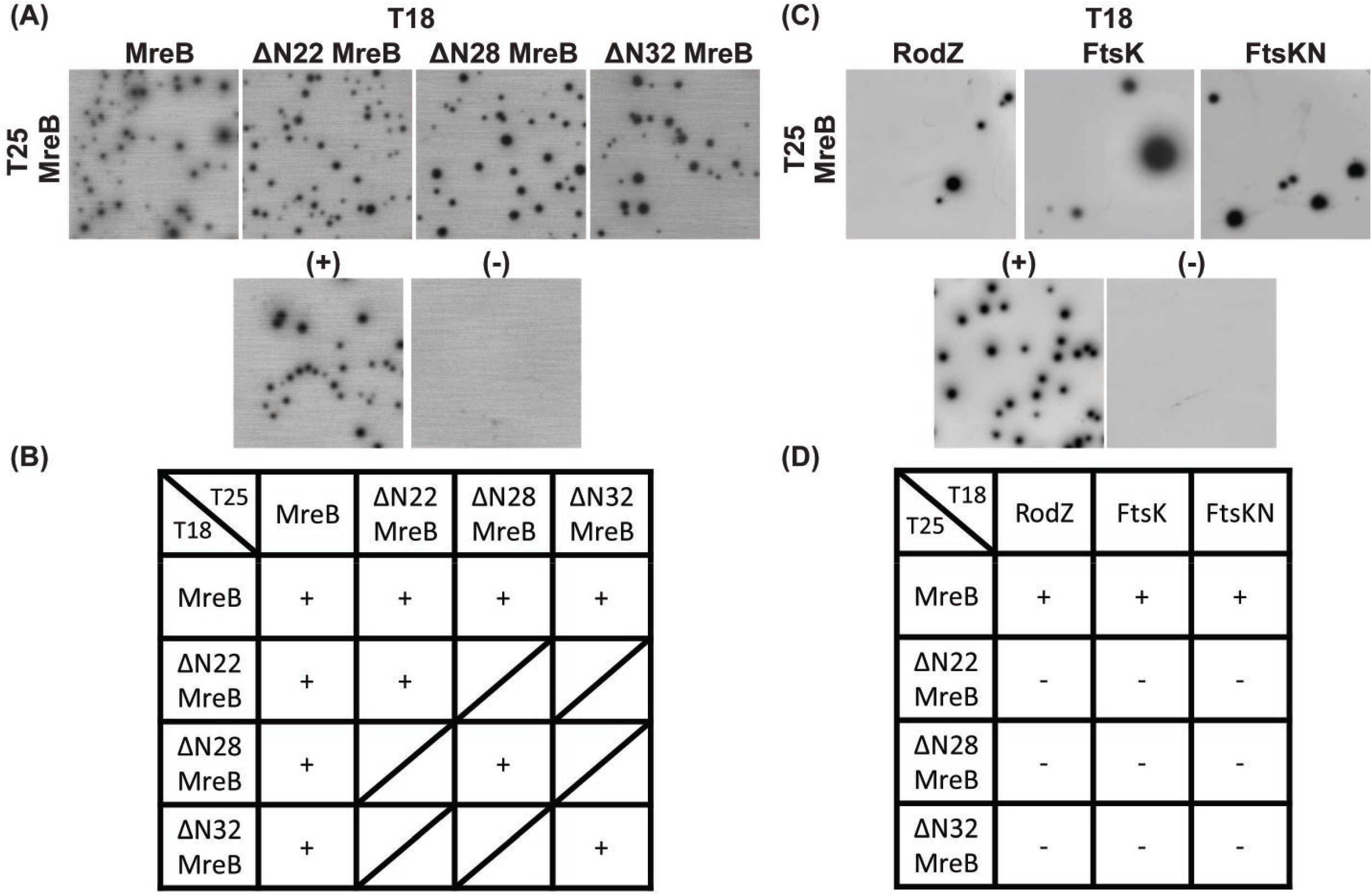
BACTH assay to test protein-protein interactions of full-length and truncated chlamydial MreB. *E. coli* DHT1 (Δ*cya*) was co-transformed with vectors encoding the indicated genes fused to the T25 and T18 catalytic domains of the *B. pertussis* adenylate cyclase. Transformants were plated on M63 minimal medium plates containing 50 μg/mL ampicillin, 25 μg/mL kanamycin, 0.5 mM IPTG, 40 μg/mL X-gal, 0.04% casamino acid, and 0.2% maltose. The plates were incubated at 30°C for 5-7 days. (A) BACTH assay of the interaction between full-length and truncated chlamydial MreBs. (B) BACTH assay of the interaction between full-length or truncated chlamydial MreBs and previously described chlamydial cell division components FtsK, the N-terminal domain of FtsK (FtsKN), and RodZ. A positive (+) control is the interaction between T25-Zip and T18-Zip, the GCN4 leucine zipper motif, and a negative (−) control is the lack of interaction between T25 and the mixture of T18-MreB and T18-ΔN22 MreB. These tests were performed a minimum of two times. ‘+’: Interaction; ‘-’: No interaction; Slash box: Not tested.

## Discussion

MreB is a well characterized rod-shape determining protein, which is conserved in most bacilli(18). When the *mreB* gene is deleted, the cell shape is changed from bacillus to coccoid in *E. coli*(18). MreB participates in organizing peptidoglycan (PG) synthesis by interacting with other proteins, such as MreC and RodZ, to direct Pbp2 and RodA activity at the membrane(19, 23). Recently, MreB has been observed as a dynamic cytoskeletal protein forming short filaments that rotate around the membrane perpendicularly to the longitudinal axis of the cell(11). This motion is driven by PG synthesis and is critical for cell elongation and maintenance of rod shape(24). Given these characteristics and properties of MreB, it is surprising that *Chlamydia*, a Gram-negative coccoid bacterium that lacks PG in its cell wall, encodes MreB. To date, we have been unable to complement any structural components of the *E. coli* divisome using chlamydial orthologs (e.g. FtsQ(6)) or Mre system (MreB (this study) or RodZ(22)), which indicates the importance of recapitulating all necessary interactions to efficiently complement. In contrast, cytosolic components of the PG synthesis pathway possessing enzymatic activity have been shown to complement *E. coli* conditional mutants(25–27). These data are consistent with the notion that the PG synthesis machinery is conserved in *Chlamydia* but the mechanisms for spatially regulating its synthesis are not conserved. These observations further suggest that the maintenance of cell morphology may not be the primary function of chlamydial MreB(5, 28). Rather, given the function of chlamydial PG strictly in cell division, chlamydial MreB may primarily function in this process.

Since *Chlamydia* lacks the conserved cell division organizing protein FtsZ, we have hypothesized and presented evidence that MreB is a functional substitute for FtsZ(6–8, 22). However, this raises the intriguing question of how MreB, which is highly conserved between *Chlamydia* and phylogenetically unrelated bacteria, can serve as the divisome organizer. Interestingly, when MreB is reintroduced into MreB-depleted cells in *B. subtilis*, a budding shape is observed that is enriched in MreB(29). This eventually leads to recruitment of cell wall machinery and formation of a rod-shape. However, in *Chlamydia*, which also undergoes a budding-like polarized division, this process is spatially restricted, thus what prevents *Chlamydia* from continuing to synthesize PG to produce a rod shape? One hypothesis is that chlamydial MreB exhibits a restricted distribution that is at least in part dependent upon its unique structural features, and we have shown here that the unique N-terminal region of chlamydial MreB can facilitate the membrane association of this cytoskeletal protein.

Initiating this study, our goal was to capture MreB localization at high resolution and at early stages of division since it becomes more difficult to resolve individual organisms within the inclusion as RBs multiply. However, we were unable to detect endogenous MreB using the previously reported antibody (13). We, therefore, took advantage of genetic tools that have been recently developed for *Chlamydia* to transform this bacterium with a plasmid construct encoding an inducible MreB_6xH. Using this strategy, we detected MreB_6xH localized at the division septum during the polarized division of the first RB. More interestingly, MreB formed a ring at the septum in the dividing cell that resembled the FtsZ-ring at the septum of *E. coli* (Fig.1). The septal MreB ring is very similar to the PG ring previously observed in *Chlamydia*(5). We speculate that the MreB puncta previously observed may be due to the low affinity of the MreB antibody as we were also unable to detect MreB at later stages during infection using this reagent. However, we also observed areas along the MreB-ring of more intense staining (Fig. 1 arrowhead). These regions may be the puncta reported previously and may represent areas of active PG synthesis. However, to date we have been unable to image both PG and MreB_6xH, perhaps due to the aldehyde fixation methods required to preserve the budding morphology of chlamydiae. Given the effects of inhibiting MreB activity on chlamydial division and PG synthesis(7, 8, 16), these data further suggest that chlamydial MreB likely has a scaffolding function for cell divisome proteins to direct PG synthesis in a manner similar to the function of FtsZ in *E. coli*(30).

Many Gram-negative MreB homologs possess an amphipathic helix at their N-terminus that allows MreB to associate with the inner membrane(9). *Chlamydia* species have conserved this feature. The most striking difference between the chlamydial species and other “canonical” MreB homologs is the presence of an extended N-terminus of 23 amino acids (aa1-23). Amphipathic prediction algorithms suggested this region possesses amphipathicity. Interestingly, canonical MreB filaments are excluded from areas of high membrane curvature in bacilli that are typically enriched in anionic phospholipids such as cardiolipin (i.e. the cell poles(31)), but *Chlamydia* are coccoid with its membrane displaying curvature similar to the poles of bacilli. Therefore, we hypothesized that the additional residues at the N-terminus of chlamydial MreB may allow it to associate more efficiently with membranes that exhibit curvature. In addition, these residues may be critical for establishing polarity in this organism as we previously observed that inhibiting MreB activity with A22 or MP265 not only blocks cell division but also depolarizes the RB(7, 8).

The amphipathic helix of canonical Gram-negative MreB homologs is critical for the association of MreB with membranes. This was demonstrated by two means: expressing two copies of the helix in tandem fused to the N-terminus of GFP and deleting the helix from MreB and assessing its localization(9). In the former, GFP was directed to the inner membrane(9). In the latter, MreB is no longer localized as membrane patches but rather as cytosolic aggregates in *E. coli*(9). Using the former approach, we observed that two copies, but not one copy, of the chlamydial amphipathic helix were sufficient to direct MreB to the membrane, consistent with what has been observed in *E. coli* (Fig. 4B). However, the additional N-terminal region of chlamydial MreB (aa1-23) together with a single copy of the conserved amphipathic region (aa24-28) directed GFP to the membrane. Intriguingly, the membrane localization was restricted to the cell polar region but did not appear to be an inclusion body since in an *mreB*-deficient mutant *E. coli*, which has a spherical morphology, the peptide fluorescence was more uniformly distributed in the membrane (Suppl. Fig. 4). Whether this region recognizes specific lipid moieties that directs its polar localization is under investigation.

We conclude that the additional N-terminal region of chlamydial MreB encodes a membrane-targeting function. Indeed, this was supported by (i) the ability of this region to direct GFP to the membrane when expressed in *Chlamydia* and (ii) the loss of GFP membrane localization when the amphipathicity of the N-terminal region was disrupted by mutagenesis. We excluded any possible effects of alternative start codons encoded by leucine residues by replicating our results using the N-terminus of *C. suis*, which does not encode any UUG codons for leucine in this region. We propose that chlamydial MreB, with both its extended N-terminus and amphipathic helix, may be more tightly associated with membranes and is critical for it to form ring structures associated with regions of high membrane curvature.

As *Chlamydia* is an obligate intracellular bacterium, it is time-consuming and difficult to genetically modify its genes in a targeted manner, and there is a high failure rate in doing so. Ideally, we would create a conditional knockout such that we could express truncated mutants of chlamydial MreB in the absence of the chromosomal full-length copy. We did attempt to localize truncation mutants of MreB in *Chlamydia*, but only the mutant lacking all predicted amphipathicity showed reduced membrane localization (Suppl. Fig. 7). Not surprisingly, we detected interactions between the MreB truncations and the full-length MreB, thus one interpretation is that the truncations, when expressed in *Chlamydia*, interacted with full-length MreB (Fig. 6), which is itself at the membrane. Furthermore, the N-terminal region is important for interactions with other proteins related to cell division and PG synthesis. When the region was deleted, chlamydial MreB no longer interacted with RodZ and FtsK by bacterial two-hybrid analysis. One possible explanation of these data is that the truncated MreB isoforms do not associate with the membrane efficiently enough to interact with the membrane proteins RodZ and FtsK. Nevertheless, these data suggest the extended N-terminal region may function as a scaffolding platform for MreB to interact with other proteins related to cell division and may be crucial for the cell divisome formation at the septum, like FtsZ.

Outstanding questions remain: how is polarity established in *Chlamydia*? Is it MreB-dependent or are other factors involved? Is MreB actively excluded from other parts of the membrane by an unknown system similar to the function of the MinCDE system in restricting FtsZ to division septa during binary fission? In this study, we performed the first systematic investigation of chlamydial MreB and its function as a cell division coordinator. Our data suggest that chlamydial MreB has multiple membrane-associating domains that may promote its assembly at the septum. This in turn may allow it to functionally substitute for the role of FtsZ in organizing the division septum.

## Materials and Methods

### Organisms and Cell Culture

McCoy (kind gift of Dr. Harlan Caldwell) and HeLa (ATCC, Manassas, VA) cells were cultured at 37°C with 5% CO_2_ in Dulbecco’s Modified Eagle Medium (DMEM; Invitrogen, Waltham, MA) containing 10% fetal bovine serum (FBS; Hyclone, Logan, UT) and 10 μg/mL gentamicin (Gibco, Waltham, MA). *Chlamydia trachomatis* serovar L2 (strain 434/Bu) lacking the endogenous plasmid (-pL2) was infected and propagated in McCoy cells for use in transformations. HeLa cells were infected with chlamydial transformants in DMEM containing 10% FBS, 10 μg/mL gentamicin, 1 U/mL penicillin G, and 1 μg/mL cycloheximide. All cell cultures and chlamydial stocks were routinely tested for *Mycoplasma* contamination using the Mycoplasma PCR detection kit (Sigma, St. Louis, MO). For *E. coli*, wild-type MG1655 and the *mreB*-deficient strain P2733 (kind gift of Dr. Piet de Boer) were cultured at 37°C with 225 rpm shaking in Lysogeny broth (LB) media containing no or several antibiotics as indicated: 50 μg/mL spectinomycin, 34 μg/mL chloramphenicol, or 25 μg/mL tetracycline. All chemicals and antibiotics were obtained from Sigma unless otherwise noted.

### Cloning

The plasmids and primers used in this study are described in Supplemental Table 1. The wildtype and truncated *Chlamydia trachomatis mreB* genes were amplified by PCR with Phusion DNA polymerase (NEB, Ipswich, MA) using 100 ng *C. trachomatis* L2 genomic DNA as a template. Some gene segments were directly synthesized as a gBlock fragment (Integrated DNA Technologies, Coralville, IA). The PCR products were purified using a PCR purification kit (Qiagen, Hilden, Germany). The HiFi Assembly reaction master mix (NEB) was used according to the manufacturer’s instructions in conjunction with plasmids pASK-GFP-mKate2-L2 (kind gift of Dr. P. Scott Hefty) cut with FastDigest AgeI and EagI (Thermofisher, Waltham, MA), pBOMB4-Tet (kind gift of Dr. Ted Hackstadt) cut with EagI and KpnI, pKT25 or pUT18C cut with BamHI and EcoRI, or pBAD33 cut with XbaI and SalI depending on the construct being prepared. All plasmids were also dephosphorylated with FastAP (ThermoFisher). The products of the HiFi reaction were transformed into either NEB-10beta (for chlamydial transformation plasmids) or DH5alpha (I^q^) competent cells (NEB), plated on appropriate antibiotics, and plasmids were subsequently isolated from individual colonies grown overnight in LB broth by using a mini-prep kit (Qiagen). For chlamydial transformation, the constructs were transformed into *dam*^-^ *dcm*^-^ competent cells (NEB) and purified as de-methylated constructs.

### Bioinformatics Analysis

Sequences for *Chlamydia trachomatis* serovar L2/434, *Pseudomonas aeruginosa* (PAO1), and *Vibrio cholerae* (O395) were obtained from the NCBI database (https://www.ncbi.nlm.nih.gov/) and for *E. coli* MG1655 from Ecocyc database (https://ecocyc.org/)(32). Protein sequence alignment was performed using Clustal Omega website (https://www.ebi.ac.uk/Tools/msa/clustalo/)(33) and the ESPript3 program (http://espript.ibcp.fr)(34). Helical wheels were made by using Helixator (http://www.tcdb.org/progs/helical_wheel.php)(35). Amphipathic helixes were predicted by using Amphipaseek program (https://npsa-prabi.ibcp.fr/cgi-bin/npsa_automat.pl?page=/NPSA/npsa_amphipaseek.html)(36).

### Transformation of *Chlamydia trachomatis*

McCoy cells were plated in a six-well plate the day before beginning the transformation procedure. *Chlamydia trachomatis* serovar L2 without plasmid (-pL2) (kind gift of Dr. Ian Clarke) was incubated with 2 μg demethylated plasmid in Tris-CaCl_2_ buffer (10 mM Tris-Cl, 50 mM CaCl2) for 30 min at room temperature. During this step, the McCoy cells were washed with 2 mL Hank’s Balanced Salt Solution (HBSS) media containing Ca^2+^ and Mg^2+^ (Gibco). After that, McCoy cells were infected with the transformants in 2 mL HBSS per well. The plate was centrifuged at 400 *x g* for 15 min at room temperature and incubated at 37°C for 15 min. The inoculum was aspirated, and DMEM containing 10% FBS and 10 μg/mL gentamicin was added per well. At 8 h post infection (hpi), the media was changed to media containing 1 μg/mL cycloheximide and 1 or 2 U/mL penicillin G, and the plate was incubated at 37°C until 48 hpi. At 48 hpi, the transformants were harvested and used to infect a new McCoy cell monolayer. These harvest and infection steps were repeated every 48 hpi until mature inclusions were observed.

### Complementation assay

The arabinose-inducible pBAD33 vectors encoding nothing, *E. coli* MreB, chlamydial MreB, or ΔN22 chlamydial MreB were transformed into both wild-type (MG1655) and *mreB* mutant strains (P2733; kind gift of Dr. Piet de Boer). Chemically competent cells of each strain were prepared using standard techniques with CaCl_2_. The strains were cultured overnight, diluted 1/50 in LB media containing chloramphenicol (34 μg/mL), tetracycline (25 μg/mL) or spectinomycin (50 μg/mL) and cultured for 2 hours when 0.01% arabinose was added as an inducer or cells were left uninduced as a control. After 2 hours and 6 hours of induction, 4 μL of the cells were mounted on 1% LB agar pad and covered with a cover slip. The morphologies of the strains were observed by a Zeiss Imager.Z2 equipped with an Apotome2 using a 100X objective.

### The localization of chlamydial MreB fusion proteins in *E. coli*

The *E. coli* MG1655 wild-type and Δ*mreB* mutant (P2733) transformed with the arabinose-inducible pBAD33G vectors encoding GFP with various MreB N-terminal peptides were cultured at 37°C with 255 rpm shaking overnight. Overnight cultures were diluted 1:100 (MG1655) or 1:50 (P2733) in LB media containing appropriate antibiotics and cultured at 37°C with 255 rpm shaking. After 2 hours of shaking, the cells were induced, or not, with 0.05% (w/v) arabinose and cultured for more 2 hours. Before mounting the samples, SynaptoRed-C2 (FM4-64; Cayman Chemical, Ann Arbor, MI) was added in the samples to a final concentration 1.5 μM to stain the membrane. After 15 min, 4 μL of each culture were placed on 1% LB agar pad and covered with coverslip. The samples were observed with a Zeiss Imager.Z2 equipped with an Apotome2 using a 100X objective.

### Indirect Immunofluorescence (IFA) Microscopy

HeLa cells were seeded in 24-well plates on coverslips at a density of 10^5^ cells per well the day before infection. Chlamydial strains expressing wild-type or truncated MreBs with a six-histidine tag at the C-terminus were used to infect HeLa cells in DMEM media containing penicillin G and cycloheximide. At 6 hpi, 10 nM anhydrotetracycline (aTc) was added. At 10.5 hpi, the coverslips of infected cells were washed with DPBS and fixed with fixing solution (3.2% formaldehyde and 0.022% glutaraldehyde in DPBS) for 2 min. The samples were then washed three times with DPBS and permeabilized with ice-cold 90% methanol for 1 min. Afterwards, the fixed cells were labeled with primary antibodies including goat anti-major outer-membrane protein (MOMP; Meridian, Memphis, TN), rabbit anti-MreBCT antibody (custom anti-peptide antibody directed against the C-terminus of *C. trachomatis* serovar L2 MreB; ThermoFisher), rabbit anti-Hsp60 (kind gift of Dr. Rick Morrison), mouse and rabbit anti-six histidine tag (Genscript, Nanjing, China and Abcam, Cambridge, UK, respectively). Secondary antibodies donkey anti-goat antibody (594), donkey anti-rabbit antibody (647), and donkey anti-mouse (488) were used to visualize the primary antibodies. The secondary antibodies were obtained from Invitrogen. Coverslips were observed by using either a Zeiss LSM 800 confocal microscope or a super-resolution SIM scope (Zeiss ELYRA PS.1).

### BACTH Assay

Competent DHT1 *E. coli*, an adenylate cyclase-deficient strain, were co-transformed with pKT25 and pUT18C vectors encoding the genes of interest or empty vectors and spread on M63 minimal media plates containing 50 μg/mL ampicillin, 25 μg/mL kanamycin, 0.5 mM IPTG, 40 μg/mL X-gal, 0.04% casamino acid, and 0.2% maltose. The plates were incubated at 30°C for 5-7 days. Blue colonies indicate positive interactions whereas no growth or small white colonies indicate no interactions.

## Acknowledgments

The authors would like to thank Dr. Lisa Rucks for critical review of the manuscript. The authors would like to thank the following individuals for providing reagents used in this study: Dr. Harlan Caldwell (National Institute of Health), Dr. Ian Clarke (University of Southampton), Dr. Piet de Boer (Case Western University), Dr. Ted Hackstadt (Rocky Mountain Labs/NIH), Dr. P. Scott Hefty (University of Kansas), and Dr. Rick Morrison (University of Arkansas for Medical Sciences). This work was supported by a grant from the National Institute for General Medical Science (R35GM124798-01) within the National Institutes of Health to SPO. The University of Nebraska Medical Center Advanced Microscopy Core Facility receives partial support from the National Institute for General Medical Science INBRE - P20 GM103427 and COBRE - P30 GM106397 grants, as well as support from the National Cancer Institute (NCI) for The Fred & Pamela Buffett Cancer Center Support Grant-P30 CA036727 and the Nebraska Research Initiative.

---

**Supplementary Figure 1. Localization of chlamydial MreB_GFPsw proteins in *C. trachomatis*.** HeLa cells were infected with *C. trachomatis* transformants containing aTc-inducible vectors encoding MreB_GFPsw proteins. At 12 hpi, expression of the GFP sandwich fusions was induced with 10 nM aTc, and the samples were fixed (3.2% Formaldehyde, 0.022% Glutaraldehyde in 1X PBS) at 16 hpi for 2 min and permeabilized with 90% methanol. The samples were stained for major outer membrane protein (MOMP; red) and GFP (green). Images were acquired on a Zeiss LSM 800 confocal microscope. Scale bar = 1 μm

**Supplementary Figure 2. Protein sequence alignment of the N-terminus of MreB from diverse *Chlamydia* phylum members.** The extended N-terminus of chlamydial MreB is conserved across *Chlamydia*. (A) The alignment was performed with Clustal Omega (https://www.ebi.ac.uk/Tools/msa/clustalo/) and represented with ESPript 3.0 (http://espript.ibcp.fr). (B) The AMPHIPASEEK prediction of amphipathicity for the *Waddlia* MreB ortholog. The blue and red residues represent the extended N-terminus and predicted amphipathic helix, respectively.

**Supplementary Figure 3. Test of complementation of chlamydial MreB in *E. coli* and interaction between chlamydial and *E. coli* MreBs by BACTH.** An *E. coli mreB*-deficient mutant (P2733) strain was transformed with an empty arabinose-inducible vector (A) or vectors encoding *E. coli* MreB (B), chlamydial MreB (C), or truncated chlamydial MreB lacking the extended N-terminal region (D). Stationary phase cultures were diluted to 1:50 in LB media containing 50 μg/mL spectinomycin, 25 μg/mL tetracycline, and 34 μg/mL chloramphenicol and cultured at 37°C with 225 rpm shaking for 2 h. The cells were then induced or not with 0.01% (w/v) arabinose. After induction, 4 μL of each culture at 2 h and 6 h were spotted under a 1% LB agar pad and covered with a coverslip. Images were acquired on a Zeiss Imager.Z2 equipped with an Apotome2 using a 100X objective. The arrows indicate the cells complemented by the induction of *E. coli* MreB. Scale bar = 2 μm. (E) BACTH assays were carried out to test interactions between chlamydial MreB and *E. coli* MreB. DHT1 *E. coli* were co-transformed with plasmids encoding the indicated fusion proteins and plated on M63 minimal medium containing 50 μg/mL ampicillin, 25 μg/mL kanamycin, 0.5 mM IPTG, 40 μg/mL X-gal, 0.04% casamino acid, and 0.2% maltose. The plates were incubated at 30°C for 5-7 days. A positive control is the interaction between T25-zip and T18-zip. A negative control is the lack of interaction between T25 and T18-chlamydial MreB and T18-*E. coli* MreB. These tests were performed a minimum of two times. (F) Western blotting was performed to test the expression of chlamydial MreB in strains used in the complementation assay depicted in (C&D). Whole cell lysates from cultures tested in the complementation assay were separated by SDS-PAGE and transferred to a PVDF membrane. The chlamydial MreB was detected with rabbit anti-MreB primary antibody and IRDye goat anti-rabbit 800CW (LI-COR, Lincoln, NE).

**Supplementary Figure 4. Localization of chlamydial N-terminal MreB-GFP fusion proteins in an *E. coli* Δ*mreB* mutant strain (P2733).** The *E. coli* Δ*mreB* mutant (P2733) was transformed with the arabinose-inducible vectors encoding GFP fused with diverse N-terminal regions of chlamydial MreB. Samples were prepared as described in the legend to Figure 4 with the membrane labeled with FM4-64. Images were acquired on a Zeiss Imager.Z2 equipped with an Apotome2 using a 100X objective. Scale bar = 2 μm.

**Supplementary Figure 5. Localization of various N-terminal regions of chlamydial MreB fused to GFP in A22-treated *C. trachomatis*.** *C. trachomatis* without plasmid (-pL2) was transformed with anhydrotetracycline (aTc)-inducible vectors encoding chlamydial MreB_1-23aa_-GFP fusion peptide (A), chlamydial MreB_1-28aa_-GFP fusion peptide (B), or chlamydial MreB_6xH (C). At 16 hpi, expression of the constructs was induced with 10 nM aTc, and at 18 hpi, 75 μM A22 was added to disrupt MreB localization. The samples were fixed at 20hpi with 1X DPBS containing 3.2% formaldehyde and 0.022% glutaraldehyde for 2 min. Afterwards, the samples were permeabilized with 90% methanol for 1 min. These samples were stained for major outer membrane protein (MOMP; red). The arrowheads show the membrane localization of the MreB_1-23aa_-GFP peptide. Images were acquired on a Zeiss Imager.Z2 equipped with an Apotome2 using a 100X objective. Scale bar = 2 μm.

**Supplementary Figure 6. Localization of the N-terminus of *C. suis* MreB-GFP fusion peptides in *C. trachomatis* and *E. coli*.** (A) The predicted amphipathicity of the N-terminus of *C. suis* (Cs) MreB. (B) A helical wheel prediction is shown. (C) The CsMreB1-23aa-GFP peptide is localized in the cytosol in *E. coli*. In contrast, the CsMreB_1-28aa_-GFP peptide is localized at the membrane at the poles of *E. coli*. These patterns are the same as those of *C. trachomatis* (see Figure 3). (D, E) HeLa cells were infected with *C. trachomatis* transformants containing aTc-inducible vectors encoding CsMre_B1-23aa_-GFP or CsMreB_1-28aa_-GFP fusion proteins. Expression of these fusion proteins was induced at 6 hpi or 16 hpi with 10 nM aTc. At 10.5 hpi or 20 hpi, the samples were fixed (3.2% Formaldehyde, 0.022% Glutaraldehyde in 1X PBS) for 2 min and permeabilized with 90% methanol (MeOH) for 1 min. These samples were stained for major outer membrane protein (MOMP; red) with GFP imaged in green. The arrowheads indicate the CsMre_B1-23aa_-GFP localized at the membrane (see also Figure 4). Images were acquired on a Zeiss LSM 800 confocal microscope with 63X objective. Scale bar = 0.5 μm (10.5 hpi) or 1 μm (20 hpi).

**Supplementary Figure 7. The localization of various truncated chlamydial MreBs in *C. trachomatis* L2.** (A) Representation of the various truncated chlamydial MreBs tested. The blue and red residues represent the extended N-terminus and predicted amphipathic helix, respectively. (B) *C. trachomatis* serovar L2 transformants containing aTc-inducible vectors encoding the truncated MreBs were used to infect HeLa cells. At 16 hpi, expression of the MreB_6xH constructs was induced with 10 nM aTc, and these samples were fixed (3.2% Formaldehyde, 0.022% Glutaraldehyde in 1X DPBS) at 20 hpi for 2 min and permeabilized with 90% methanol. The samples were stained for major outer membrane protein (MOMP; red) and six histidine tag (green). Images were acquired on a Zeiss Imager.Z2 equipped with an Apotome2 using a 100X objective. The white box represents the cells which are zoomed in at the upper right. Scale bar = 2 μm.

**Supplementary Table. List of Plasmids, Strains, and Primers Used in the Study.**

## Notes

#### Summary of Updates

New data in Figure 5 demonstrating that mutating the N-terminal region of MreB to reduce its amphipathicity prevents it from allowing GFP to direct to the membrane in Chlamydia. New Supplemental Figure 5 showing that treatment with A22 does not redirect the N-terminal MreB regions fused to GFP from the membrane to the cytosol. New Supplemental Figure 6 showing that the N-terminal region of C. suis behaves the same as the N-terminal region of C. trachomatis.

