## Supplemental Table 1 for "Critical Role for the Unique N-Terminus of Chlamydial MreB in Directing Its Membrane Association and Interaction with Elements of the Divisome"

**Supplementary Table 1. List of Plasmids, Strains, and Primers Used in the Study**

| **Construct Plasmid** | **Relevant genotype** | **Ori** | **Source of Reference** |
| --- | --- | --- | --- |
| pASK-GFP-mKate-L2 (pTLR2) | *bla* P*tet*::*gfp* | ColE1 | (1) |
| pTLR2-*mreB*_6xH | *bla* P*tet*::*Ctr_mreB*_6xH | ColE1 | This study |
| pTLR2-*mreB_67*_6xH | *bla* P*tet*::ΔN66nt *Ctr_mreB*_6xH | ColE1 | This study |
| pTLR2-*mreB_85*_6xH | *bla* P*tet*::ΔN84nt *Ctr_mreB*_6xH | ColE1 | This study |
| pTLR2-*mreB_97*_6xH | *bla* P*tet*::ΔN96nt *Ctr_mreB*_6xH | ColE1 | This study |
| pTLR2-*mreB*-*gfp*_sw | *bla* P*tet*::*Ctr_mreB-gfp_sw* | ColE1 | This study |
| pTLR2-*mreB_67*-*gfp*_sw | *bla* P*tet*::ΔN66nt *Ctr_mreB-gfp_sw* | ColE1 | This study |
| pTLR2-*mreB_85*-*gfp*_sw | *bla* P*tet*::ΔN84nt *Ctr_mreB-gfp_sw* | ColE1 | This study |
| pTLR2-*mreB_97*-*gfp*_sw | *bla* P*tet*::ΔN96nt *Ctr_mreB-gfp_sw* | ColE1 | This study |
| pBAD33 | *cat* P*ara* | p15A | (2) |
| pBAD33-*Ec*_*mreB* | *cat* P*ara*::*Ec_mreB* | p15A | This study |
| pBAD33-*mreB* | *cat* P*ara*::*Ctr_mreB* | p15A | This study |
| pBAD33-*mreB_*67 | *cat* P*ara*::ΔN66nt *Ctr_mreB* | p15A | This study |
| pBAD33G | *cat* P*ara*::*gfp* | p15A | This study |
| pBAD33G-*mreB*1-96nt | *cat* P*ara*::*Ctr_mreB*1-96nt*-gfp* | p15A | This study |
| pBAD33G-*mreB*1-84nt | *cat* P*ara*::*Ctr_mreB*1-84nt*-gfp* | p15A | This study |
| pBAD33G-*mreB*1-69nt | *cat* P*ara*::*Ctr_mreB*1-69nt*-gfp* | p15A | This study |
| pBAD33G-*mreB*1-69nt duplicate | *cat* P*ara*::*Ctr_mreB*1-69nt *Ctr_mreB*1-69nt *-gfp* | p15A | This study |
| pBAD33G-*mreB*67-96nt | *cat* P*ara*::*Ctr_mreB*67-96nt*-gfp* | p15A | This study |
| pBAD33G-*mreB*67-96nt duplicate | *cat* P*ara*::*Ctr_mreB*67-96nt *Ctr_mreB*67-96nt*-gfp* | p15A | This study |
| pBAD33G-*Ec_mreB* | *cat* P*ara*::*Ec_mreB-gfp* | p15A | This study |
| pBAD33G-*Ec_mreB_*22 | *cat* P*ara*::ΔN21nt *Ec_mreB-gfp* | p15A | This study |
| pBAD33G-*mreB*1-84nt-*Ec_mreB_*22 | *cat* P*ara*::*Ctr_mreB*1-84nt-ΔN21nt *Ec_mreB-gfp* | p15A | This study |
| pBAD33G-*mreB*1-69nt-*Ec_mreB_*22 | *cat* P*ara*::*Ctr_mreB*1-69nt-ΔN21nt *Ec_mreB-gfp* | p15A | This study |
| pBAD33G-L7K, L22R *mreB_1-69nt_* | *cat* P*ara*::L7K, L22R *Ctr_mreB_1-69nt_-gfp* | p15A | This study |
| pBAD33G-L7K, L22R, F25K *mreB_1-69nt_* | *cat* P*ara*::L7K, L22R, F25K *Ctr_mreB_1-84nt_-gfp* | p15A | This study |
| pBAD33G-Cs*mreB_1-69nt_* | *cat* P*ara*::*C. suis_mreB_1-69nt_-gfp* | p15A | This study |
| pBAD33G-Cs*mreB_1-84nt_* | *cat* P*ara*::*C. suis_mreB_1-84nt_-gfp* | p15A | This study |
| pBAD33G-L7K, L22R *mreB_1-69nt-_EcmreB_22* | *cat* P*ara*::L7K, L22R *Ctr_mreB_1-69nt_-* ΔN21nt *Ec_mreB-gfp* | p15A | This study |
| pBAD33G-L7K, L22R, F25K *mreB_1-84nt_* | *cat* P*ara*::L7K, L22R, F25K *Ctr_mreB_1-84nt_-* ΔN21nt *Ec_mreB-gfp* | p15A | This study |
| pBOMB4-Tet | *bla* P*tet*::*mCherry* P*Nmen*::*gfp* | pUC19 | (3) |
| pBOMB-G-Tet | *bla* P*tet*::*mCherry* | pUC19 | This study |
| pBOMB-G-Tet-*mreB*1-69nt-*gfp* | *bla* P*tet*::*Ctr_mreB*1-69nt*-gfp* | pUC19 | This study |
| pBOMB-G-Tet-*mreB*1-84nt-*gfp* | *bla* P*tet*::*Ctr_mreB*1-84nt*-gfp* | pUC19 | This study |
| pBOMB-G-Tet-L7K, L22R *mreB*1-69nt-*gfp* | *bla* P*tet*::L7K, L22R *Ctr_mreB*1-69nt*-gfp* | pUC19 | This study |
| pBOMB-G-Tet-L7K, L22R *mreB*1-84nt-*gfp* | *bla* P*tet*::L7K, L22R, F25K *Ctr_mreB*1-84nt*-gfp* | pUC19 | This study |
| pBOMB-G-Tet-Cs*mreB*1-69nt-*gfp* | *bla* P*tet*::*C. suis_mreB*1-69nt*-gfp* | pUC19 | This study |
| pBOMB-G-Tet-Cs*mreB*1-84nt-*gfp* | *bla* P*tet*::*C. suis_mreB*1-84nt*-gfp* | pUC19 | This study |
| pKT25 | *aph* P*lac*::*t25* | p15A | (4) |
| pKT25-*mreB* | *aph* P*lac*::*t25-Ctr_mreB* | p15A | (5) |
| pKT25-*mreB_*67 | *aph* P*lac*::*t25-*ΔN66nt *Ctr_mreB* | p15A | This study |
| pKT25-*mreB_*85 | *aph* P*lac*::*t25-*ΔN84nt *Ctr_mreB* | p15A | This study |
| pKT25-*mreB_*97 | *aph* P*lac*::*t25-*ΔN96nt *Ctr_mreB* | p15A | This study |
| pKT25-*Ec_mreB* | *aph* P*lac*::*t25-Ec_mreB_SW_* | p15A | (6) |
| pKT25-zip | *aph* P*lac*::*t25-zip* | p15A | (4) |
| pUT18C | *bla* P*lac*::*t18* | ColE1 | (4) |
| pUT18C-*mreB* | *bla* P*lac*::*t18-Ctr_mreB* | ColE1 | (5) |
| pUT18C-*mreB_*67 | *bla* P*lac*::*t18-*ΔN66nt *Ctr_mreB* | ColE1 | This study |
| pUT18C-*mreB_*85 | *bla* P*lac*::*t18-*ΔN84nt *Ctr_mreB* | ColE1 | This study |
| pUT18C-*mreB_*97 | *bla* P*lac*::*t18-*ΔN96nt *Ctr_mreB* | ColE1 | This study |
| pUT18C-*rodZ* | *bla* P*lac*::*t18-Ctr_rodZ* | ColE1 | (7) |
| pUT18C-*ftsK* | *bla* P*lac*::*t18-Ctr_ftsK* | ColE1 | (5) |
| pUT18C-*ftsK*N | *bla* P*lac*::*t18-*N-terminus of *Ctr_ftsK* | ColE1 | (5) |
| pUT18C-*Ec_mreB* | *bla lacI^q^* P*lac*::*t18-Ec_mreB_SW_* | ColE1 | (6) |
| pUT18C-zip | *bla* P*lac*::*t18-zip* | ColE1 | (4) |
| pFB112 | *tet sdiA* | ColE1 | (8) |
| pFB124 | *aadA* cI857(Ts) PλR::*mreC mreD*-LE | pSC101 | (8) |

| ***E. coli* Strain** | **Relevant genotype** | **Source of Reference** |
| --- | --- | --- |
| MG1655 | F^-^, lambda^-^, *rph-1* | Lab strain |
| FB17 | *dadR trpE trpA tna mreBCD<> frt* | (8) |
| DHT1 | F^-^ *glnV44* (AS) *recA1 endA1 gyrA96* (Nal^R^) *thi-1 hsdR17 spoT1 rfbD1 cya-854 ilv-691 ::Tn10 (TetR)* | (9) |
| P2733 | FB17 pFB112 pFB124 | (8) |

| **Primer name** | **Sequence** | **Features** | **Usage** |
| --- | --- | --- | --- |
| mreB 5'/pTLR2/LIC_F | tttgtttaactttaagaaggagata ATGAGCCCATACCGCAGC | lower case for plasmid overlap construction | for amplification of *mreB* into pTLR2 |
| mreB6XH/pTLR2/R | ccatttttcacttcacaggtcaacc ttaatggtgatggtgatggtg TACTAAACTCTCTTTTCGTTTCTTCAATTG | lower case for plasmid overlap construction, adds 6xH to *mreB* sequence | for amplification of *mreB* into pTLR2 |
| mreB_1_5'/pBAD33/F | ttcgagctcggtacccggggatcct tttgtttaactttaagaaggagatataca ATGAGCCCATACCGCAGC | lower case for plasmid overlap construction, adds Shine-Dalgano (SD) sequence | for amplification of *mreB* into pBAD33 |
| mreB_67_5'/pBAD33/F | ttcgagctcggtacccggggatcct tttgtttaactttaagaaggagatatacaatg GGTCGTTTCGATCGTGTATTTAATTTTTTTTC | lower case for plasmid overlap construction, adds Shine-Dalgano (SD) sequence | for amplification of ΔN66nt *mreB* into pBAD33 |
| mreB 3'/pBAD33/LIC_R | ctagaggatccccgggtaccgagct TCATACTAAACTCTCTTTTCGTTTC | lower case for plasmid overlap construction | for amplification of *mreB*s into pBAD33 |
| EcmreB/pBAD33/F | gctcggtacccggggatcct tttgtttaactttaagaaggagatataca ATGTTGAAAAAATTTCGTG | lower case for plasmid overlap construction, adds Shine-Dalgano (SD) sequence | for amplification of *Ec*_*mreB* into pBAD33 |
| EcmreB/pBAD33/R | caagcttgcatgcctgcagg TTACTCTTCGCTGAACAGG | lower case for plasmid overlap construction | for amplification of *Ec*_*mreB* into pBAD33 |
| mreB/pBAD33G/F | tgggctagcgaattcgagct tttgtttaactttaagaaggagatataca ATGAGCCCATACCGCAGC | lower case for plasmid overlap construction, adds Shine-Dalgano (SD) sequence | for amplification of *mreB_1-96nt* into pBAD33G |
| mreB1-96nt/pBAD33G/R | gtcgactctagaggatcccc c AAAAAAATTAAATACACGATCGAAACG | lower case for plasmid overlap construction | for amplification of *mreB_1-96nt* into pBAD33G |
| Ctr L2/mreB/pBAD/5' | TATATGGTACCTGATTAACTTTATAAGGAGGAAAAACAT**A**TGAGCCCATACCGCAGCTTATAT | Underline for KpnI site | for amplification of *mreB_*1-69nt and 1-84nt into pBAD vectors |
| Ctr L2/mreB_69/pBAD/3' | CGCCCTCTAGA**A**CCCAACGCCTTGTTATACAGACGGTTAGA | Underline for XbaI site | for amplification of *mreB_*1-69nt into pBAD vectors |
| Ctr L2/mreB_84/pBAD/3' | CGCCCTCTAGA**T**ACACGATCGAAACGACCCAACGCCTTGTTATAC | Underline for XbaI site | for amplification of *mreB_*1-84nt into pBAD vectors |
| mreB_67-84D/pBAD33/F | gctcggtacccggggatcct tttgtttaactttaagaaggagatataca ATGGGTCGTTTCGATCGTG | lower case for plasmid overlap construction, adds Shine-Dalgano (SD) sequence | for amplification of *mreB_67*-84nt-gfp from gBlock |
| GFP/pBAD33/R | caagcttgcatgcctgcagg TTATTTGTATAGTTCATCCATGCCATG | lower case for plasmid overlap construction | for amplification of *mreB_67*-84nt-gfp from gBlock |
| Ec mreB/pBAD/5' | CCCCCGAGCTCTGATTAACTTTATAAGGAGGAAAAACAT**A**TGCTGAAAAAATTTCGTGGCATGT | Underline for SacI site | for amplification of Ec_*mreB* into pBAD33 |
| Ec mreB_22/pBAD/5' | CCCCCGAGCTCTGATTAACTTTATAAGGAGGAAAAACAT**A**TGTTCTCCAATGACTTGTCCATTGA | Underline for SacI site | for amplification of ΔN21nt Ec_*mreB* into pBAD33 |
| Ec mreB/pBAD33G/3' | ATAATGTCGAC**C**TCTTCGCTGAACAGGTCGCCGCCGTGCATGTCGATC | Underline for SalI site | for amplification of Ec_*mreB* into pBAD33G |
| mreB/pBOMB/F | gatctaaagaggagaaaggatctgc ATGAGCCCATACCGCAG | lower case for plasmid overlap construction | for amplification of *mreB_*1-69nt and 1-84nt fused with *gfp* into pBOMB-G-Tet |
| GFP/pBOMB/R | tttgaatggtcgaccggtac TTATTTGTATAGTTCATCCATGCCATG | lower case for plasmid overlap construction | for amplification of *mreB_*1-69nt and 1-84nt fused with *gfp* into pBOMB-G-Tet |
| mreB_4/T25/F | ctgcagggtcgactctagag AGCCCATACCGCAGCTTATATAAG | lower case for plasmid overlap construction | for amplification of *mreB* into pKT25 |
| mreB_67/T25/F | ctgcagggtcgactctagag GGTCGTTTCGATCGTGTATTTAATTTTTTTTC | lower case for plasmid overlap construction | for amplification of ΔN66nt *mreB* into pKT25 |
| mreB_85/T25/F | ctgcagggtcgactctagag TTTAATTTTTTTTCCGGG | lower case for plasmid overlap construction | for amplification of ΔN84nt *mreB* into pKT25 |
| mreB_97/T25/F | ctgcagggtcgactctagag TCCGGGAATGTTGGTATC | lower case for plasmid overlap construction | for amplification of ΔN96nt *mreB* into pKT25 |
| mreB_1101/(pKT25)/R | tcacgacgttgtaaaacgacggccg TCATACTAAACTCTCTTTTCGTTTC | lower case for plasmid overlap construction | for amplification of *mreBs* into pKT25 |
| mreB_4/T18/F | actgcaggtcgactctagag AGCCCATACCGCAGCTTATATAAG | lower case for plasmid overlap construction | for amplification of *mreB* into pUT18C |
| mreB_67/T18/F | actgcaggtcgactctagag GGTCGTTTCGATCGTGTATTTAATTTTTTTTC | lower case for plasmid overlap construction | for amplification of ΔN66nt *mreB* into pUT18C |
| mreB_85/T18/F | actgcaggtcgactctagag TTTAATTTTTTTTCCGGG | lower case for plasmid overlap construction | for amplification of ΔN84nt *mreB* into pUT18C |
| mreB_97/T18/F | actgcaggtcgactctagag TCCGGGAATGTTGGTATC | lower case for plasmid overlap construction | for amplification of ΔN96nt *mreB* into pUT18C |
| mreB_1101/(pUT18C)/R | accatattacttagttatatcgatg TCATACTAAACTCTCTTTTCGTTTC | lower case for plasmid overlap construction | for amplification of *mreBs* into pUT18C |
| Ct009/rodZ/BACTH/5'BamHI | ATATAGGATCCA**G**GTAGAGCAAACCAGGGGAATTAC | Underline for BamHI site | for amplification of *rodZ* |
| Ct009/rodZ/BACTH/3'KpnI | AGGGTGGTACC**T**TA**G**AAAAGGTTGAATAGATTCCCTAG | Underline for KpnI site | for amplification of *rodZ* |
| Ct739/ftsK/BACTH/5’XbaI | ACTAGCTGGTTCTAGAT**G**GAAAAGAACGGAAGAAA | Underline for XbaI site | for amplification of *ct739*(*Ctr_ftsK*) |
| Ct739/ftsK/BACTH/3’KpnI | CACAGCTAGAGGTACCAT**T**TAATCGTCCTGATTTGA | Underline for KpnI site | for amplification of *ct739*(*Ctr_ftsK*) |
| Ct739/ftsKN/BACTH/5'XbaI | ACTAGTCTAGAT**G**GAAAAGAACGGAAGAAA | Underline for XbaI site | for amplification of N-terminus of *ct739*(*Ctr_ftsK*) |
| Ct739/ftsKN/BACTH/3'KpnI | CTAGAGGTACCAT**T**TGAGAAATTTTAGGAGA | Underline for KpnI site | for amplification of N-terminus of *ct739*(*Ctr_ftsK*) |
| mreB67 5'/pTLR2/LIC_F | tttgtttaactttaagaaggagata ATGGGTCGTTTCGATCGTG | lower case for plasmid overlap construction | for amplification of ΔN66nt *mreB* into pTLR2 |
| mreB85 5'/pTLR2/LIC_F | tttgtttaactttaagaaggagata ATGTTTAATTTTTTTTCCGGGAATG | lower case for plasmid overlap construction | for amplification of ΔN84nt *mreB* into pTLR2 |
| mreB_97_5'/pTLR2/F | tttgtttaactttaagaaggagata atgTCCGGGAATGTTGGTATC | lower case for plasmid overlap construction | for amplification of ΔN96nt *mreB* into pTLR2 |
| gfp/pTLR2-mreB/LIC F | ttatccgttaggtggaggtagt AGTAAAGGAGAAGCACTTTTC | lower case for plasmid overlap construction | for amplification of *gfp* into pTLR2 |
| gfp/pTLR2-mreB/LIC R | cctgatcactacctcc GAGTCCGGACTTGTATAGTTC | lower case for *mreB* 3' insert DNA overlap construction | for amplification of *gfp* into pTLR2 |
| mreB 5'/pTLR2/LIC_R2 | cctttactactacctcc ACCTAACGGATAAGCAGAACCTATAG | lower case for *gfp* insert DNA overlap construction | for amplification of N-terminus of *mreB* into pTLR2 |
| mreB 3'/pTLR2/LIC_F2 | caagtccggactc *ggaggtagt* GATCAGGAATTGGAGATG | lower case for *gfp* insert DNA overlap construction, adds link residues, GGS | for amplification of C-terminus of *mreB* into pTLR2 |
| Ec mreB/BACTH/5'Bam | CCTTGGGATCCT**C**TGAAAAAATTTCGTGGCATGT | Underline for BamHI site | for amplification of Ec_*mreB* |
| Ec mreB/BACTH/3'Sacstp | ATATTGAGCTC**T**TA**C**TCTTCGCTGAACAGGTCGCC | Underline for SacI site | for amplification of Ec_*mreB* |
| Ec mreB/BACTH/3'Sac | TATATGAGCTCCC**C**TCTTCGCTGAACAGGTCGCC | Underline for SacI site | for amplification of Ec_*mreB* |
