## Supplementary Figures for "Critical Role for the Unique N-Terminus of Chlamydial MreB in Directing Its Membrane Association and Interaction with Elements of the Divisome"

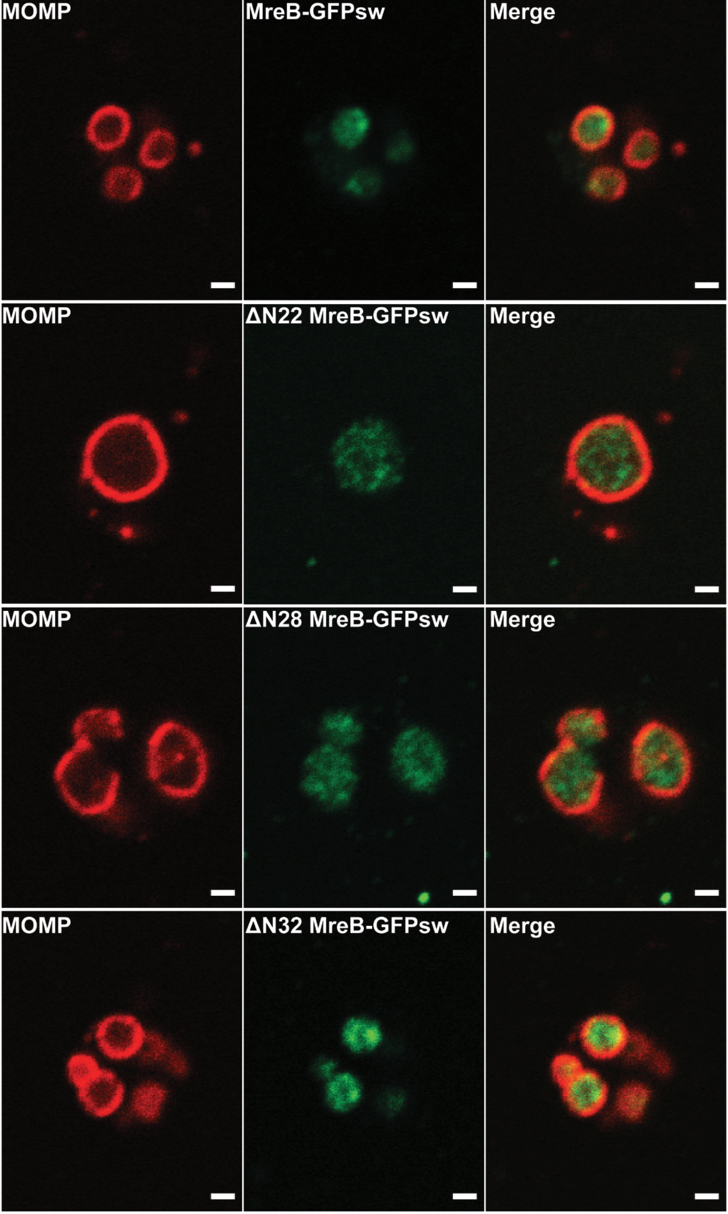


Supplemental Figure 1

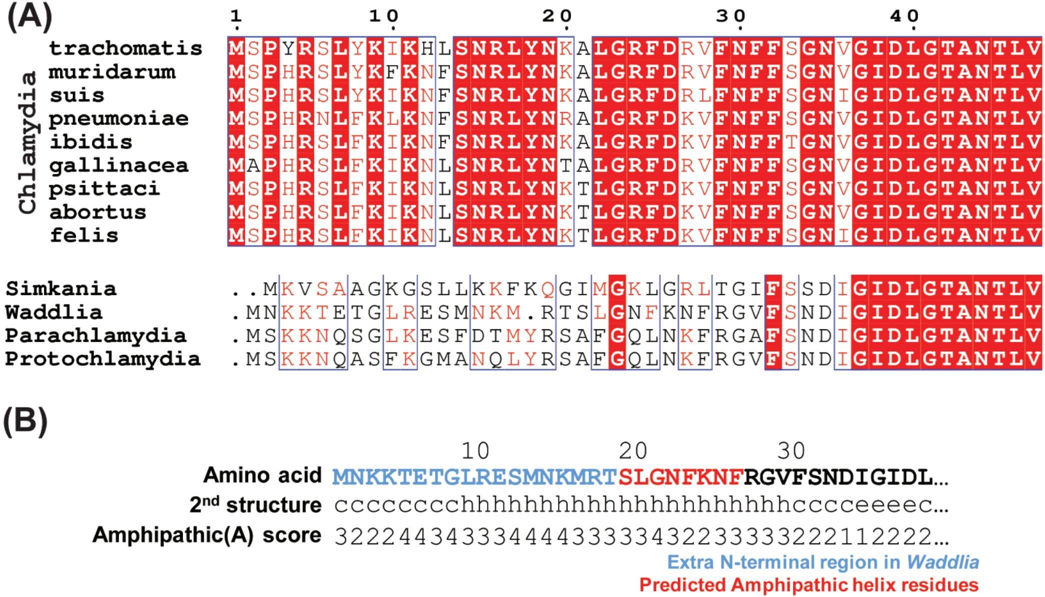


Supplemental Figure 2

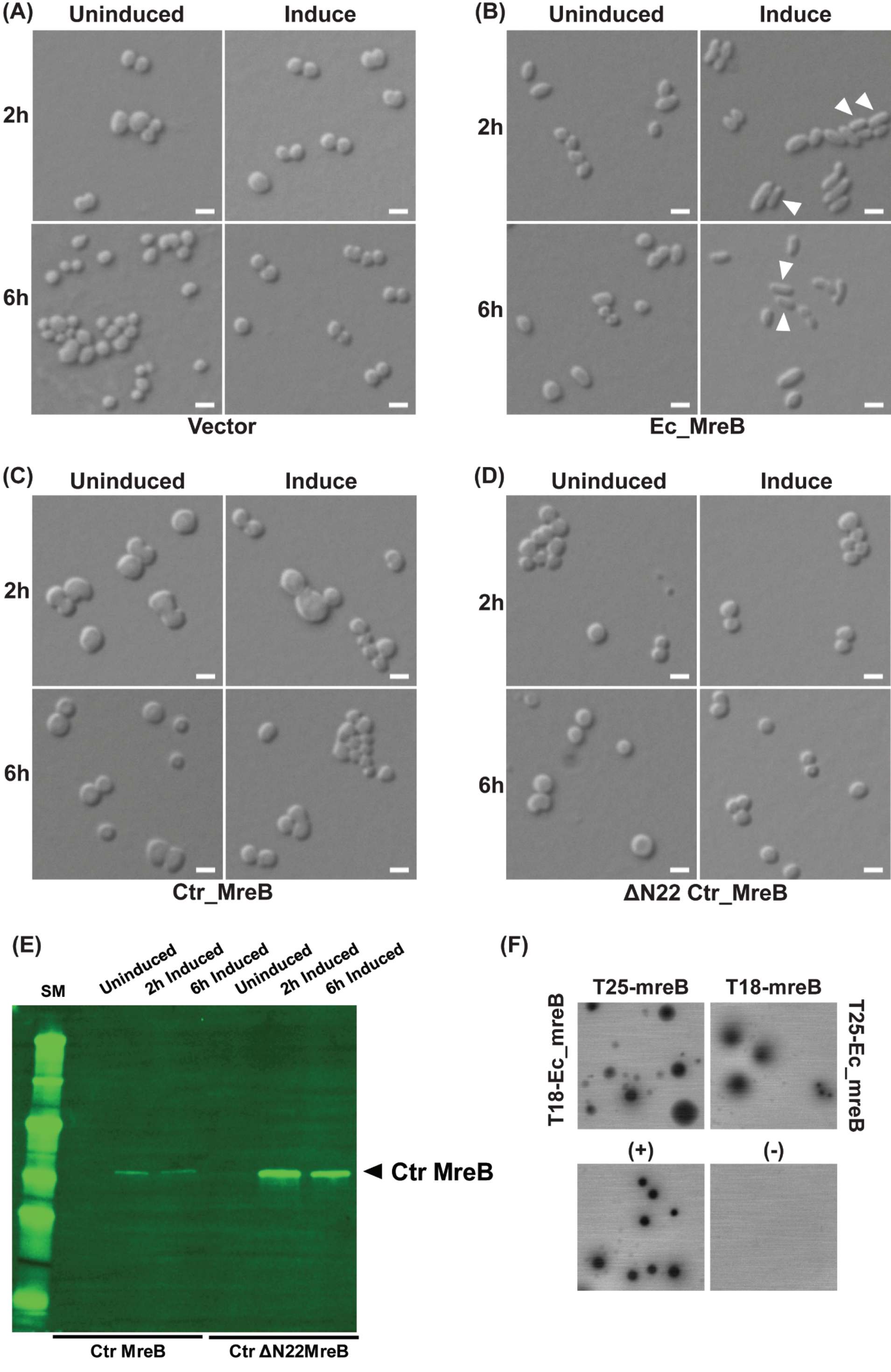


Supplemental Figure 3

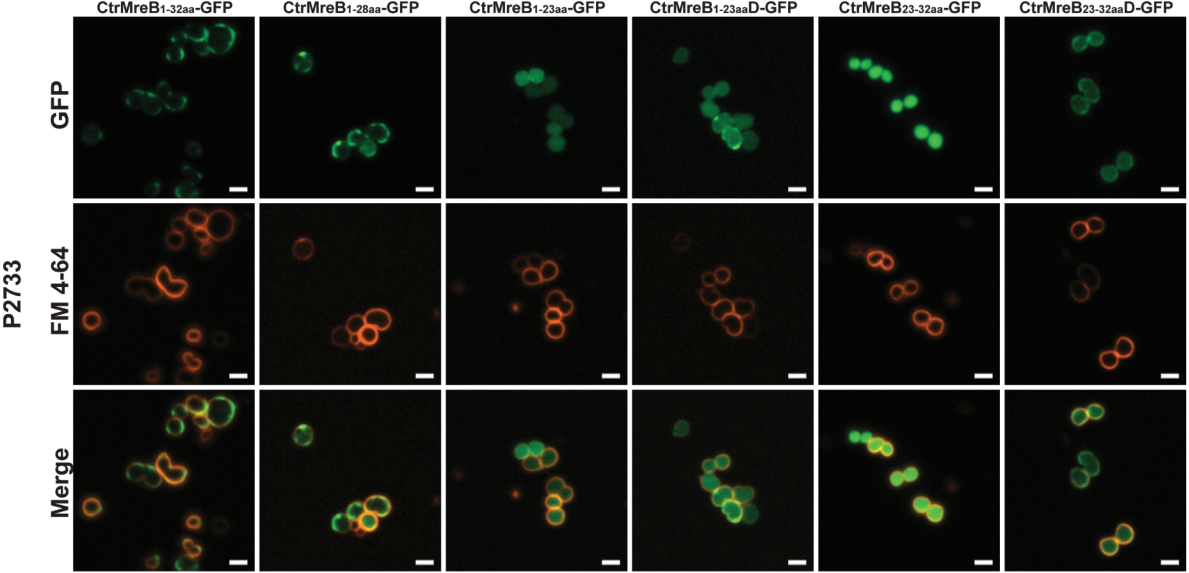


Supplemental Figure 4

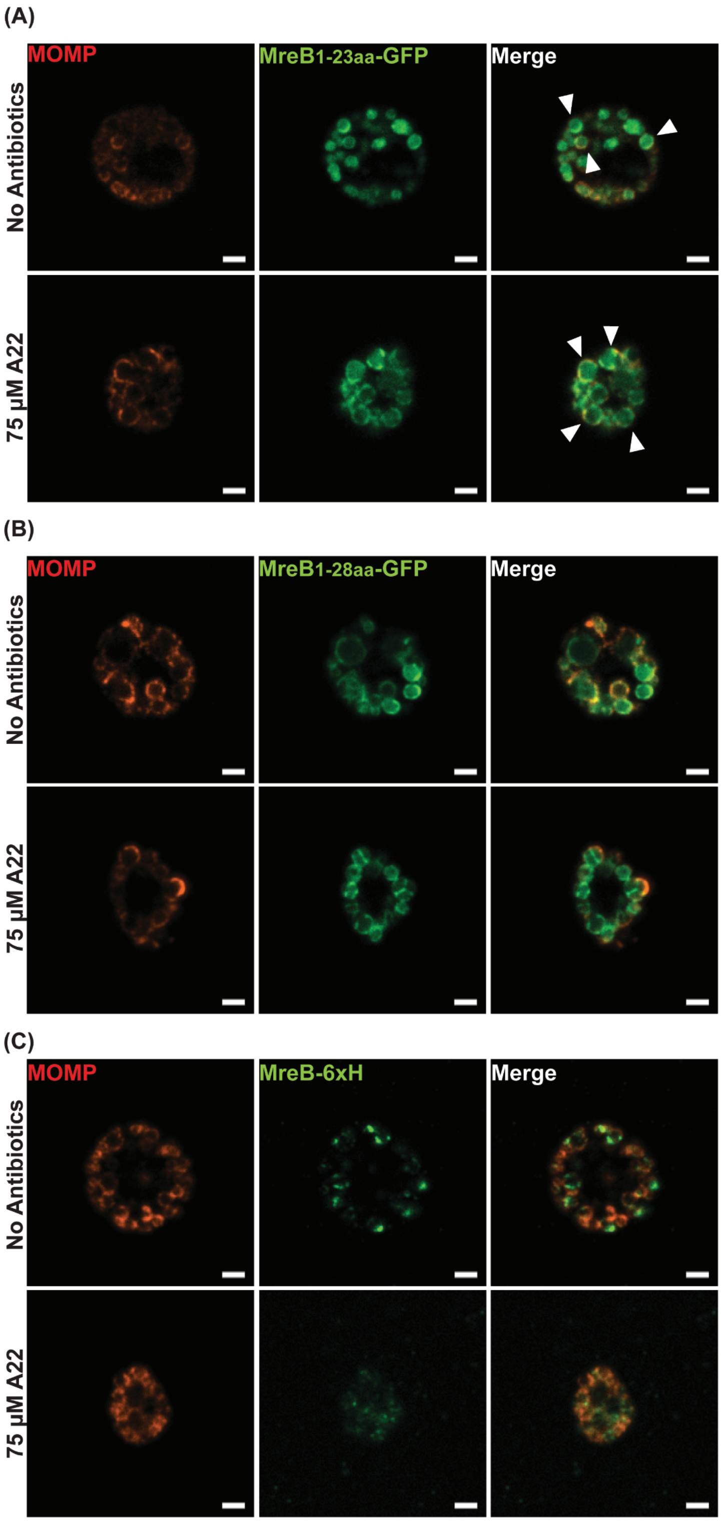


Supplemental Figure 5

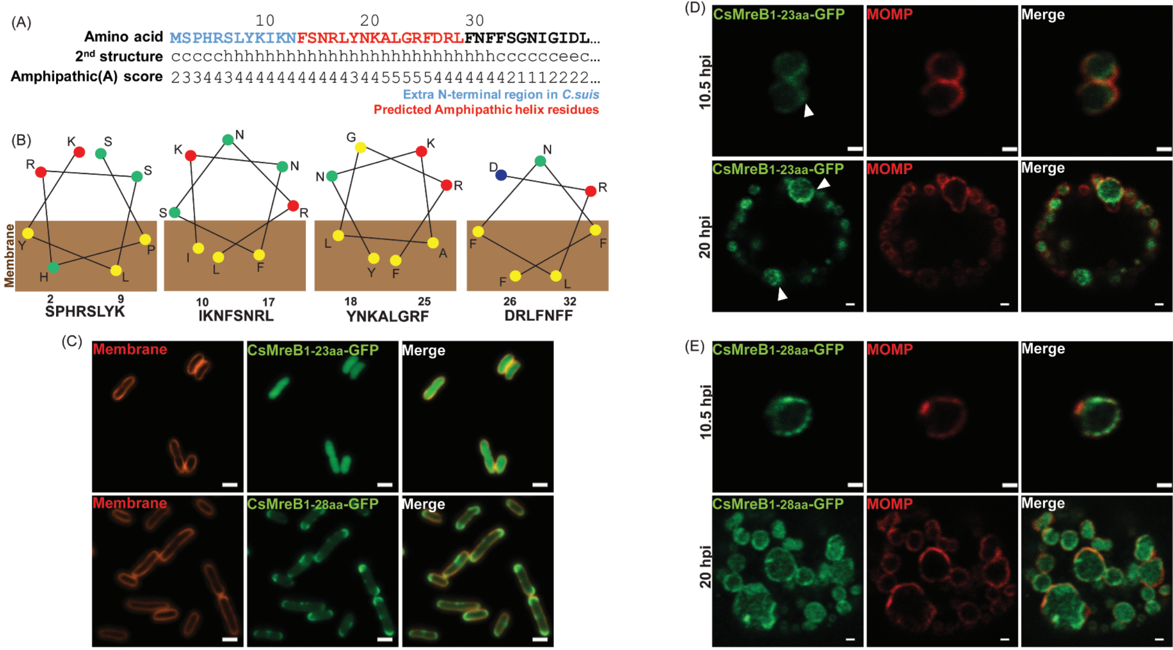


Supplemental Figure 6

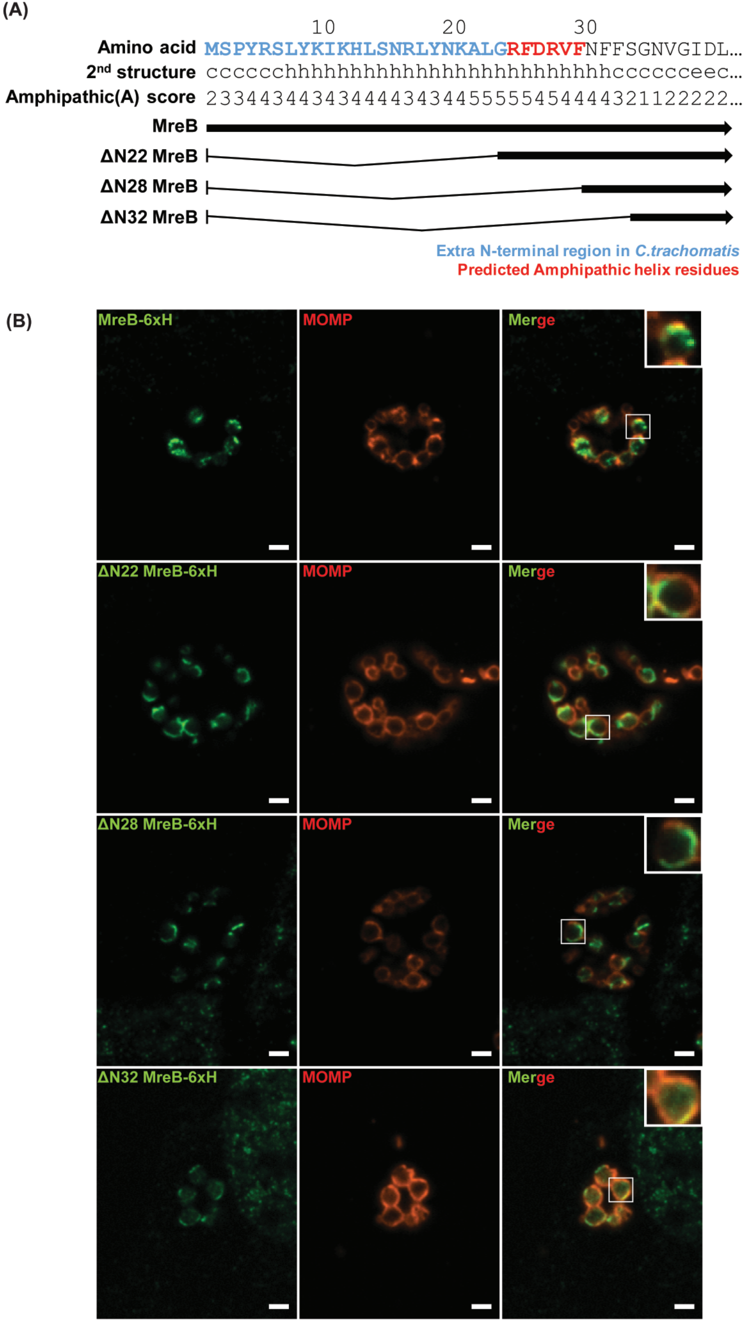


Supplemental Figure 7
